# Metabolite mimicry identifies butyrate analogs with select protective functions in the intestinal mucosa

**DOI:** 10.64898/2026.01.07.697087

**Authors:** Alfredo Ornelas, Jacob A. Countess, Ji Yeon Kim, Rachel H. Cohen, Brittany D. Gomez, Rebecca L. Roer, Faiz Minhajuddin, Kiranmayee Yenugudhati Vijaya Sai, Liheng Zhou, Julia L. M. Dunn, Philip Reigan, Ian M. Cartwright, Caroline H. T. Hall, Geetha Bhagavatula, Joseph C. Onyiah, Alexander S. Dowdell, Sean P. Colgan

**Affiliations:** Mucosal Inflammation Program and Division of Gastroenterology and Hepatology; Medical Scientist Training Program, Department of Medicine, University of Colorado School of Medicine; Division of Gastroenterology, Hepatology, and Nutrition, Department of Pediatrics, University of Colorado School of Medicine; Department of Pharmaceutical Sciences, Skaggs School of Pharmacy and Pharmaceutical Sciences, University of Colorado Anschutz Medical Campus, Aurora CO, 80045; Children’s Hospital Colorado, Aurora, CO 80045; Department of Medicine, Rocky Mountain Veterans Regional VA Medical Center, Aurora, CO 80045

**Keywords:** metabolite-mimicry, butyrate, colitis, wound healing, intestinal-barrier, butyrate

## Abstract

Microbial-derived short-chain fatty acids regulate a variety of pathways in the healthy colonic mucosa. In particular, butyrate serves as the primary energy source for colonocytes and regulates gene transcription by stabilizing the transcription factor hypoxia-inducible-factors (HIF) and functioning as a histone deacetylase (HDAC) inhibitor. A limitation of butyrate as a therapeutic is its rapid metabolism in differentiated colonocytes. Furthermore, intestinal stem cells (ISCs) respond differently to butyrate, preferentially using glucose for energy procurement. To address these limitations, we explored metabolite-mimicry to discover compounds with potent or selective biological responses within the butyrate pathway(s). We discovered an analog, 3-chlorobutyrate (3-Cl BA), that significantly enhances epithelial barrier formation and wound healing in vitro. Mechanistically, we revealed that 3-Cl BA is a potent HDAC inhibitor. Furthermore, unlike butyrate, 3-Cl BA does not stabilize HIF and it is not used as metabolic fuel. In vivo studies in a DSS-colitis model revealed that contrary to butyrate, 3-Cl BA is protective. Studies in stem-like colonoids demonstrated that only butyrate inhibits ISC proliferation and differentiation. Furthermore, it was recently reported that HIF stabilization inhibits ISCs activity. Given the fact that butyrate but not 3-Cl BA stabilizes HIF, we surmised that 3-Cl BA would circumvent these detrimental functional consequences. We demonstrate here that pharmacologic HIF stabilization inhibits colonoid differentiation and that genetic loss of HIF significantly promotes ISC differentiation. This study reveals a promising butyrate analog protective in colitis and demonstrates the advantages of metabolite-mimicry to dissect selective biological functions from major metabolites in the gut.

**Significance statement:** Butyrate is a well-studied microbial short-chain fatty acid that regulates a number of mucosal pathways and is paramount in maintaining intestinal integrity. In health, it is a major source of energy for colonocytes and regulates gene transcription. The role of butyrate in disease is still controversial and not well understood. When butyrate is not metabolized or well-utilized (e.g. disease), it accumulates in intestinal stem cells leading to reduced cell proliferation and differentiation, thereby hampering intestinal barrier recovery. In this study, we describe a butyrate analog that enhances epithelial barrier formation and wound healing. Furthermore, as opposed to native butyrate, this butyrate analog is protective in a colitis mouse model and does not exhibit detrimental influences on intestinal stem cells.

Graphical abstract
Microbially derived butyrate plays a key role in intestinal homeostasis. It is the primary source of energy for colonocytes, contributing to a metabolic and oxygen gradient as it is metabolized by differentiated cells along the intestinal crypt. Through the regulation of transcription factors such as HIF and the inhibition of HDAC, it regulates barrier formation and wound healing promoting a strong tight junction profile. Furthermore, well oxygenated ISCs at the bottom of the crypt are unaccustomed to the effects of butyrate, including HIF stabilization (left). In disease, loss of intestinal architecture leads to a disrupted metabolic/oxygen gradient where butyrate accumulates in stem cells leading to decreased proliferation, differentiation, and increases in apoptosis. 3-Cl BA selectively acts as an HDACi and does not stabilize HIF, exhibiting no significant detrimental effects on ISCs (right). Created in BioRender. Ornelas, A. (2026) https://BioRender.com/x9cy8mw

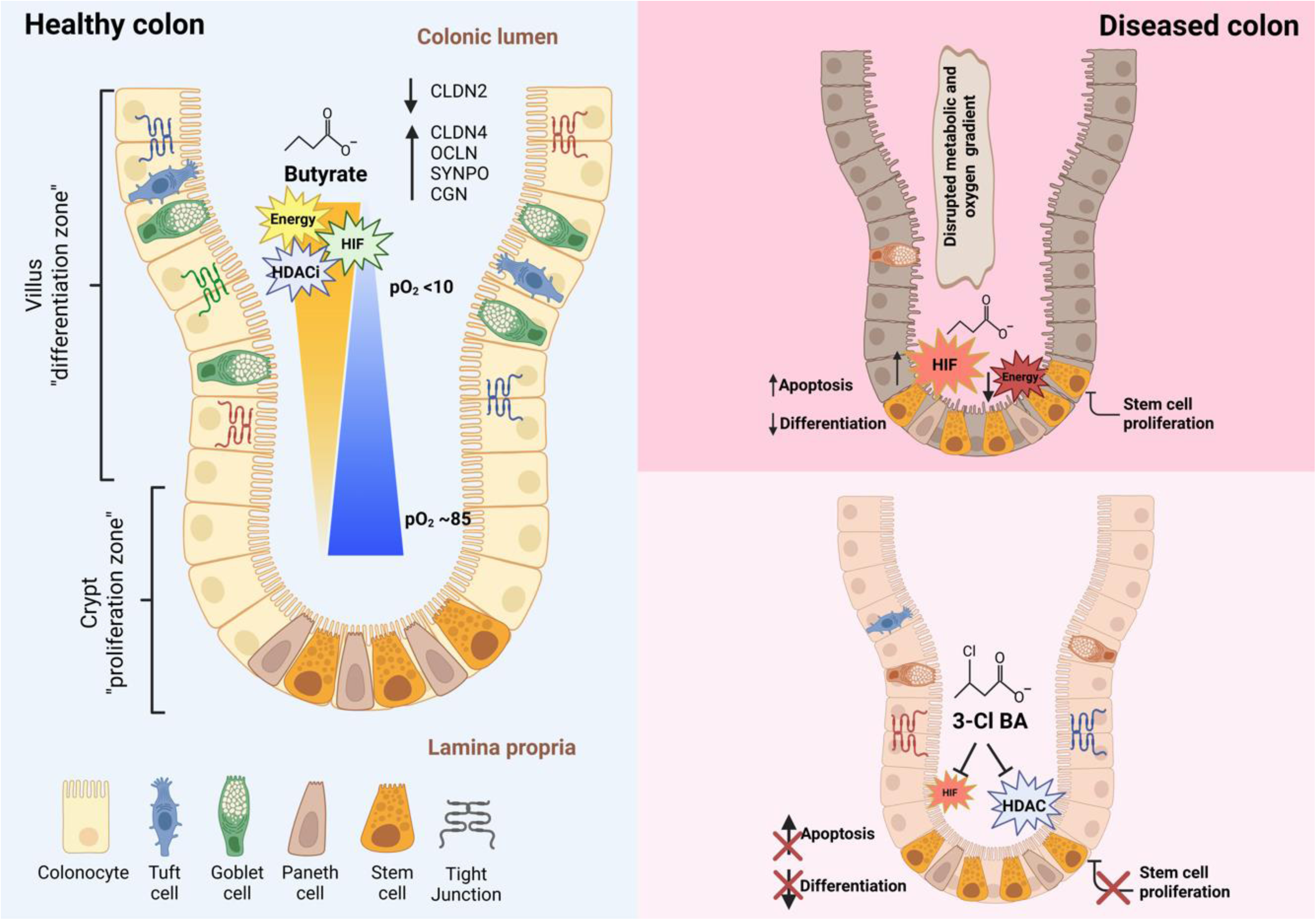

## Introduction

Termed “gut microbiota”, the gastrointestinal tract is home to trillions of microorganisms, where over a thousand diverse bacterial species dominate the microbial population and play a key role in various biological processes in health and disease.^1,2^ Thought to exist in a 1:1 ratio of human and bacterial cells,^3^ the microbiota contributes over 150 times more genetic information than that of the entire human genome.^4^ Known as the “hidden organ”, the gut microbiota is a primary mediator of homeostasis, regulating nutrient metabolism, barrier integrity, inflammation, defending against pathogens, and immune cell signaling.^5^

The gut microbiota is responsible for carrying out the vital task of producing beneficial metabolites such as short-chain fatty acids (SCFAs), indole derivatives, amine-containing compounds, secondary bile acids, and vitamins. SCFA are the products of bacterial fermentation of insoluble fiber, primarily in the colon^6^ and play key beneficial roles to maintain gut health.^7^ SCFA are fatty acids containing 1-6 carbon atoms and can display various functional groups including amines, alcohols, or branched carbon chains. The three primary SCFAs produced in the intestine are acetate (∼60%), propionate (∼20%), and butyrate (∼20%). Depending on dietary substrates, SCFAs can reach tolerable high concentrations in the colon of up to 150 mM.^8–10^ SCFAs are well known for their role in colon energy supply, gut barrier regulation, influence on immune responses, and playing a signaling role in the gut-brain axis.^11–14^

Of the SCFA, butyrate has been most widely studied for its benefits to the host. Butyrate is involved in cell cycle progression, amelioration of mucosal inflammation, improving wound healing, epithelial barrier formation, and as an antioxidant.^15^ As the preferred energy source in the colon, up to 95% of butyrate is absorbed by colonocytes and transformed into fuel via β-oxidation and the tricarboxylic acid cycle.^16,17^ The remaining butyrate is capable of orchestrating the regulation of a variety of biological pathways to maintain intestinal barrier homeostasis: Butyrate is a ligand for G-protein coupled receptors^18–20^ and it has been reported that butyrate either directly or indirectly activates the aryl hydrocarbon receptor transcription factor.^21,22^ Furthermore, our laboratory originally reported that butyrate stabilizes the transcription factor hypoxia-inducible factor (HIF), a master transcriptional regulator of many genes involved in intestinal homeostasis.^23,24^ Lastly, it is also well known that butyrate is a potent HDAC inhibitor (HDACi) ^25–27^, making butyrate one of the more important metabolites generated by the gut microbiota.

It is well documented that patients with inflammatory bowel disease (IBD) exhibit significantly decreased levels of butyrate and butyrate-producing bacteria, termed dysbiosis.^28^ As butyrate is mostly used as an energy source, such dysbiosis potentially limits its clinical application in activating other biological pathways, especially when butyrate levels are low. Furthermore, not all intestinal epithelial cells (IECs) metabolize butyrate equally. For example, it has long been known that IEC from patients with IBD show deficient utilization of butyrate in the conversion to energy, the so-called “starved gut hypothesis”^29^. Moreover, it has been shown that undifferentiated colonocytes, such as intestinal stem cells (ISCs) at the base of the crypt, prefer glucose over butyrate as an energy source and thus are susceptible to the downstream effects of increased butyrate levels.^30–32^

Metabolite-mimicry, defined as the creation of synthetic molecules that mimic biological activity of naturally occurring metabolites, is a relatively new and understudied field that can potentially accelerate drug discovery.^33–37^ In theory, biomimicry can lead to potent drug-like molecules similar to naturally occurring metabolites that regulate specific biological pathways. Given the broad biological actions of butyrate in the gut, our laboratory has focused on discovering butyrate-mimicking compounds that exhibit more specificity or potency towards various biological pathways. In these efforts, we previously reported a compound that stabilizes HIF more potently and in a prolonged manner compared to the parent compound.^33^ In the present studies, we discovered a butyrate-mimicking compound, 3-chlorobutyrate (3-Cl BA), that significantly enhanced IEC barrier formation, promoted epithelial wound healing, functioned as a potent HDACi but possessed no HIF stabilization activity. Unlike butyrate, 3-Cl BA showed significant anti-inflammatory properties in a mouse model of colitis. These results contribute to the discovery of tolerable metabolite-mimicking drugs that may alleviate mucosal inflammation in diseases such as IBD.

## Results

### 3-Cl BA enhances epithelial barrier function and accelerates IEC wound healing similar to butyrate

Butyrate is well known to enhance intestinal epithelial barrier formation and accelerate wound healing, presumably functioning as an energy source for colonocytes. Based on these studies, we profiled a small library of structurally related butyrate compounds on IEC barrier formation and wound healing (a representative panel of butyrate-mimicking compounds is shown in Fig. 1*A*). First, we studied epithelial barrier formation by monitoring transepithelial electrical resistance (TEERs), as we have described previously.^33,38^ For these purposes, T84 (Fig. 1*B*) and Caco-2 cells (SI Appendix Fig. 1) were grown on semipermeable membranes overnight followed by exposure to various butyrate derivatives (all sodium salts at 5 mM) or PBS. Monolayers were exposed to compounds on both the basolateral and apical surfaces and TEERs values were monitored every 24 h. Both butyrate and 3-Cl BA significantly promoted epithelial barrier formation, with noticeable differences as early as 24 h post-treatment and up to a four-fold increase in TEER after 48 h (Fig 1*B*). Interestingly, other closely related molecules including the structural isomer 4-chloro butyrate (4-Cl BA) and a potential hydrolysis byproduct (3-hydroxy butyrate [3-OH BA]) had no influence in epithelial barrier formation beyond that of vehicle alone. We also evaluated functional barrier activity through fluorescein isothiocyanate-dextran (FITC-dextran) paracellular flux assays, as we have described in the past.^38^ T84 cells were exposed to butyrate and 3-Cl BA (5 mM) and grown to maximum TEER reading prior to FITC-dextran flux assay performed every 30 min for a total of 120 min. By this measure of barrier function, we observed that butyrate and 3-chlorobutyrate significantly enhanced barrier as measured by paracellular flux compared to vehicle (Fig 1*C*).

**Figure 1.**
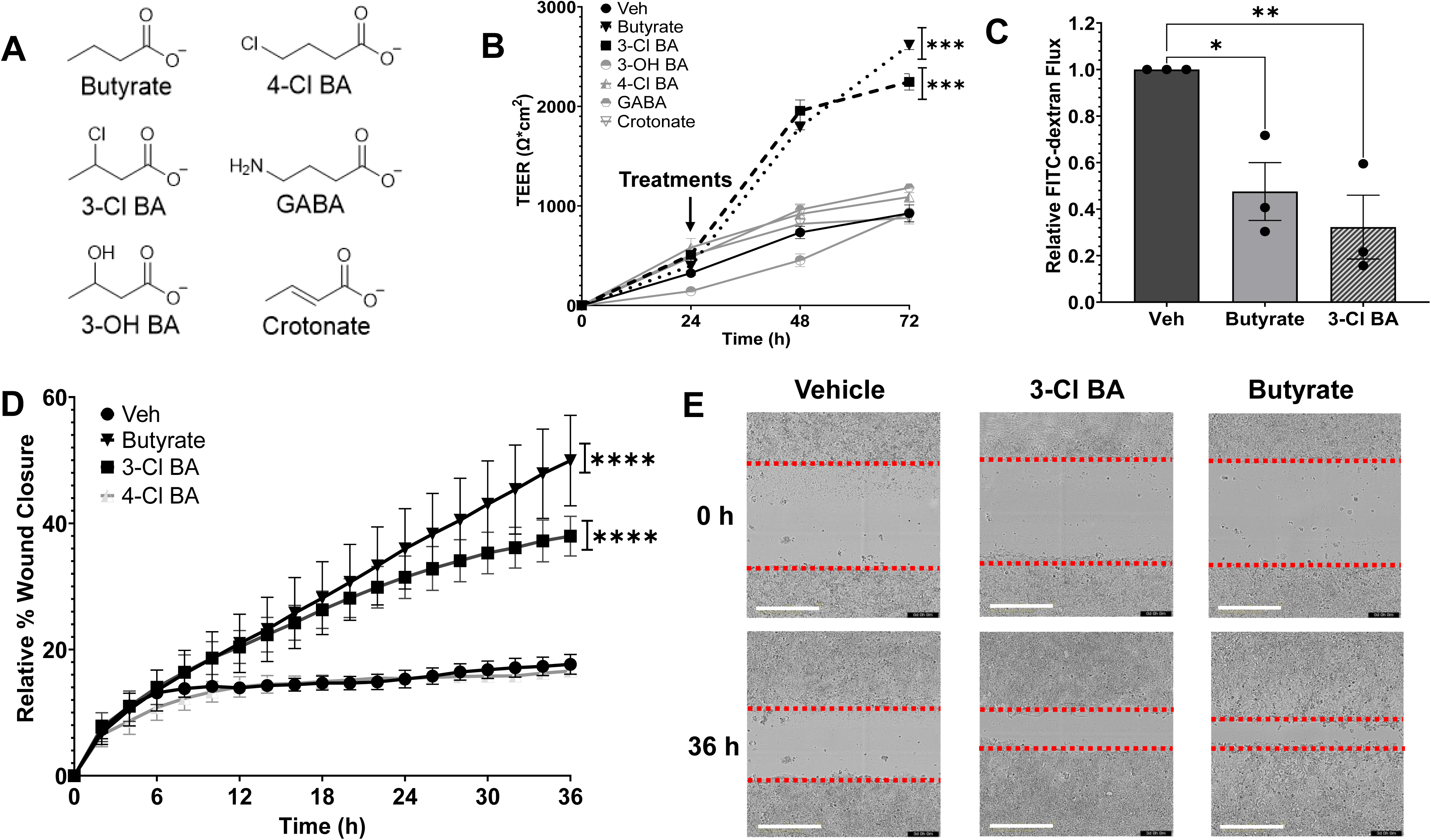
Influence of 3-Cl BA in barrier formation, permeability, and wound healing. (A) Representative panel of selected screened butyrate-mimicking compounds. (B) Epithelial barrier formation over time in monolayers of T84 cells exposed to various butyrate-mimicking derivatives (all at 5 mM, Veh = PBS). (*n = 3-4;* error bars: SEM, \*\**p* < 0.01, \*\*\**p* < 0.001, \*\*\*\**p* < 0.0001 by Two-way ANOVA with the Geisser-Greenhouse correction, Dunnett’s multiple comparisons test). (C) Cell layer permeability flux rate assay using 4-kDa FITC-dextran in T84 cells treated with 5 mM butyrate, 3-Cl BA or PBS. Data presented as relative FITC-dextran flux normalized to vehicle treated cells. (*n = 3;* error bars: SEM, \**p* < 0.05, \*\**p* < 0.01 by One-way ANOVA, Fisher’s multiple comparison). (D) Scratch wound healing monitored over time by relative wound closure percentage in T84 cells treated with butyrate, 3-Cl BA, 4-Cl BA (all at 5 mM) or PBS. (*n = 3;* error bars: SEM, \*\*\*\**p* < 0.0001 by Two-way ANOVA with the Geisser-Greenhouse correction, Dunnett’s multiple comparisons test). (E) Live cell images of scratch wound healing (0 h and 36 h) in T84 cells vehicle (PBS) control cells and cells treated with 5 mM butyrate or 3-Cl BA (red dotted line, cell migration/wound edge. Scale bar: 400 μm).

IEC proliferation and migration demand significant energy expenditure. We reasoned that butyrate, being a preferred colonocyte energy source, would significantly accelerate wound healing compared to other butyrate mimicking compounds. We explored wound healing through scratch wound assays. Briefly, T84 cells were grown to confluence in 96-well plates, uniformly scratch wounded, and wound healing was monitored by image capture every 2 h for 36 h. The images obtained were evaluated for relative wound closure percentage over time. As shown in Figure 1*D*, butyrate and 3-Cl BA significantly promoted wound healing compared to vehicle controls. Of note, the closely related structural isomer 4-Cl BA did not accelerate wound healing beyond that of controls. Figure 1*E* depicts representative images of the original wound and the striking wound closure in monolayers of IECs exposed to both butyrate and 3-Cl BA at 36 h.

We assessed potential effects 3-Cl BA might exert on IECs by measuring proliferation and potential cytotoxicity. We discovered that butyrate but not 3-Cl BA significantly reduces cell proliferation in T84 cells (SI Appendix Fig. 2*A*). Cytotoxicity studies in T84 cells revealed that both the native butyrate and 3-Cl BA exhibit very mild toxicity levels, although, 3-Cl BA (∼5%) was significantly less toxic than butyrate (∼15%) in T84 cells (SI Appendix Fig. 2*B*). Lastly, to establish the stability of 3-Cl BA in aqueous conditions, nuclear magnetic resonance (NMR) studies were performed by dissolving 3-Cl BA in deuterated water and comparing original ^1^H and ^13^C NMR spectra (0 h) to spectra after 24 h incubation at 37 °C. NMR studies revealed there is no change to the structural integrity of 3-Cl BA including no observed hydrolysis (SI Appendix Fig.3-6)

**Figure 2.**
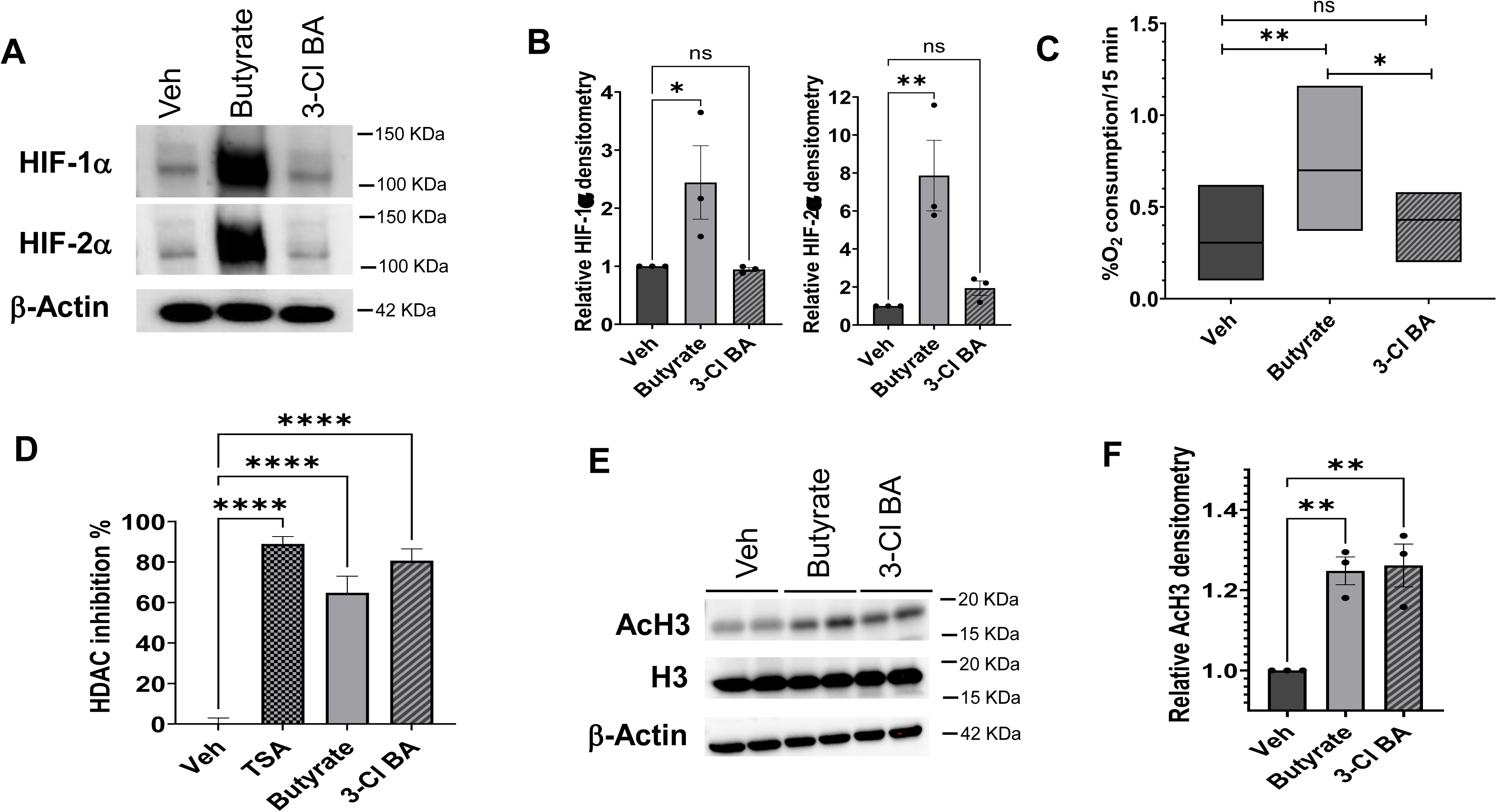
3-Cl BA does not stabilize HIF or consume oxygen in IECs, but it is a potent HDAC inhibitor. (A) Representative immunoblots of HIF-1α and HIF-2α protein levels in T84 cells exposed to 5 mM butyrate, 3-Cl BA, or PBS for 6 h in normal oxygenation conditions. (B) HIF-1α and HIF-2α were quantified using actin-normalized densitometry. (*n = 3*; error bars: SEM, \**p* < 0.05, \*\**p* < 0.01 by One-way ANOVA, Fisher’s multiple comparison). (C) Rates of oxygen consumption were calculated from linear regression of oxygen saturation data in Caco-2 monolayers on inserts treated with 5 mM butyrate, 3-Cl BA or PBS; (*n = 9* inserts from 3 independent experiments, presented as floating bars with line at median, and analyzed by ordinary One-way ANOVA, Fisher’s multiple comparison, \**p* < 0.05, \*\**p* < 0.01.) (D) Percentage HDAC inhibition in nuclear extracts of Caco-2 cells exposed to butyrate or 3-Cl BA (5 mM), Veh = PBS, or positive control = TSA at 1 μM. (*n = 4*; error bars: SEM, \*\*\*\**p* < 0.0001 by One-way ANOVA, Fisher’s multiple comparison). (E) Immunoblot of acetylated histone H3, Histone H3, and actin protein levels in Caco-2 cells exposed to butyrate or 3-Cl BA (5 mM, 24h) and (F) quantification of acetylation of histone-3 (AcH3) by actin-normalized densitometry of both H3 and Ac-H3, followed by relative densitometry of Ac-H3 to H3. (*n = 3*; error bars: SEM, \*\**p* < 0.01 by One-way ANOVA, Fisher’s multiple comparison).

**Figure 3.**
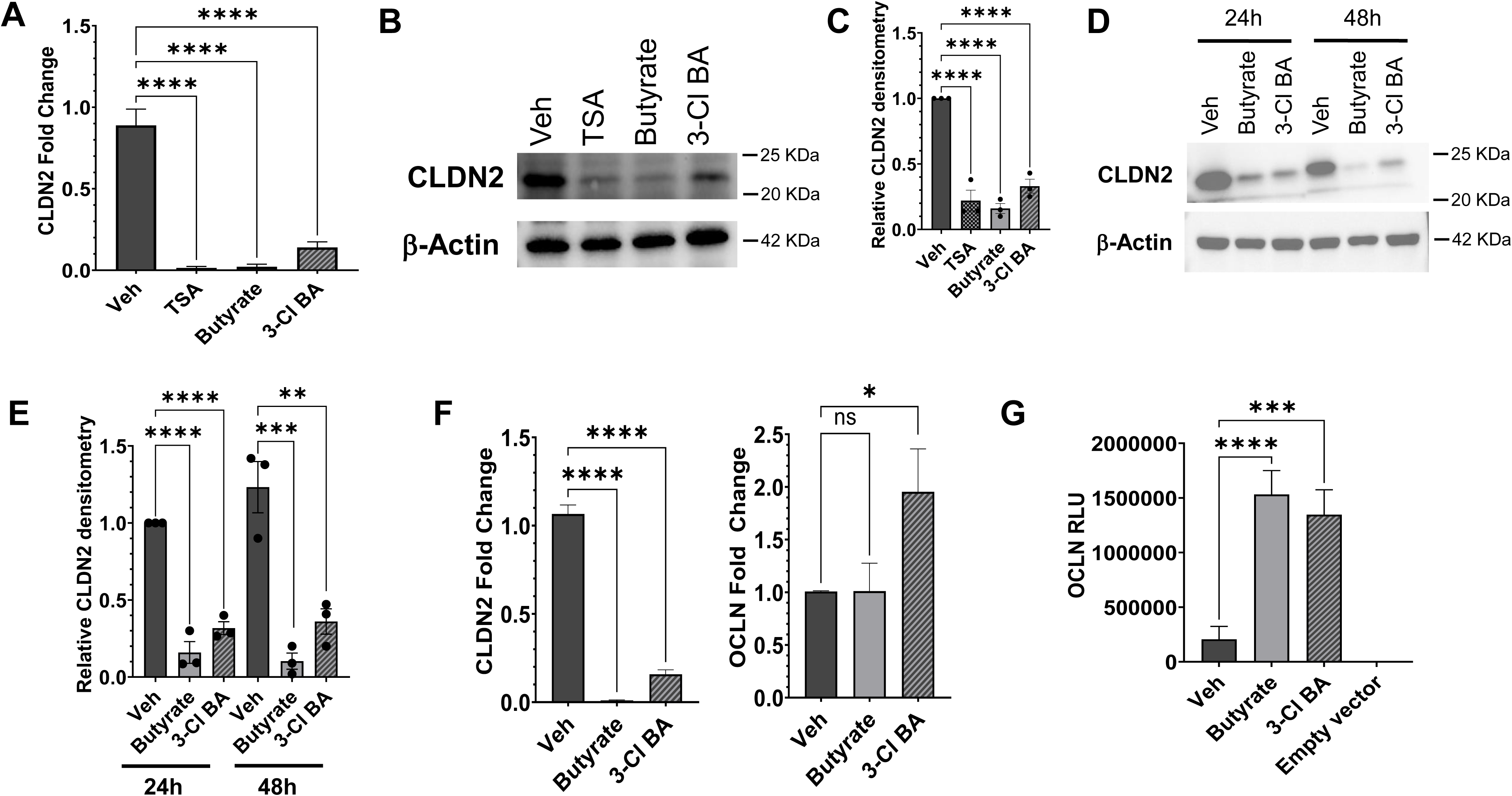
Butyrate and 3-Cl BA potently downregulate the “leaky claudin” CLDN2 and upregulate OCLN in IECs. (A) *CLDN2* mRNA expression and (B) CLND2 protein levels in Caco-2 cells treated with 5 mM butyrate or 3-Cl BA. TSA used as positive control at 1 μM and Veh = PBS. Cells were exposed to treatments for 18 h. (mRNA expression *n = 3*; error bars: SEM, \*\*\*\**p* < 0.0001 by One-way ANOVA, Fisher’s multiple comparison). (C) Quantification of CLDN2 protein was performed using actin-normalized densitometry. (*n = 3*; error bars: SEM, \*\*\*\**p* < 0.0001 by One-way ANOVA, Fisher’s multiple comparison). (D) CLDN2 protein levels in timecourse immunoblot of Caco-2 cells exposed to butyrate, 3-Cl BA (5 mM) or Veh = PBS for 24 or 48 h. (E) Quantification of CLDN2 protein in timecourse experiment was performed using actin-normalized densitometry. (*n = 3*; error bars: SEM, \*\**p* < 0.01, \*\*\**p* < 0.001, \*\*\*\**p* < 0.0001 by One-way ANOVA, Fisher’s multiple comparison). (F) *CLDN2* and *OCLN* mRNA expression in T84 cells treated with 5 mM butyrate, 3-Cl BA or Veh = PBS for 18 h. (*n = 3*; error bars: SEM, \**p* < 0.05, \*\*\*\**p* < 0.0001 by One-way ANOVA, Fisher’s multiple comparison). (G) HeLa cells were transfected with an empty vector control and an OCLN promoter reporter (OCLN-luc). After 24 h, cells were treated with PBS, 5 mM butyrate or 3-Cl BA for another 24 h and luciferase activity was quantified. (*n = 3*; error bars: SEM, \*\*\**p* < 0.001, \*\*\*\**p* < 0.0001 by One-way ANOVA, Fisher’s multiple comparison).

### 3-Cl BA exhibits selective biological roles acting as a potent HDAC inhibitor

We next focused on distinguishing mechanistic endpoints between native butyrate and the 3-Cl BA mimetic. Here, we profiled HIF stabilization, oxygen consumption and HDAC inhibition. As shown in Figure 2*A* and 2*B*, western blot analysis revealed that butyrate but not 3-Cl BA (both at 5 mM, 6 h, normoxia) stabilized HIF-1α and HIF-2α protein in T84 cells. Next, we extended our studies to assess the influence of butyrate and 3-Cl BA in O_2_ consumption which we have shown serves as a surrogate measure of oxidative phosphorylation^23,24^. Using Oxo-dish assays,^23,24^ we monitored oxygen consumption in Caco-2 monolayers plated on semipermeable membranes and exposed to PBS, butyrate (5 mM), or 3-Cl BA (5 mM). As shown in Fig. 2*C*, the rate of O_2_ consumption was significantly increased by butyrate over vehicle and 3-Cl BA, suggesting that 3-Cl BA is not used as fuel for energy procurement by IEC. Finally, we compared butyrate and 3-Cl BA function as inhibitors of several classes of HDACs.^39^ Using HDAC activity inhibition assays, we measured the inhibition of these enzymes in nuclear extracts of Caco-2 cells exposed to butyrate, 3-Cl BA (5 mM, 24 h) or the *bona fide* HDACi trichostatin A (TSA, 1 μM, 24 h). Indeed, as observed in Fig 2*D* we discovered that like butyrate and the HDACi TSA, 3-Cl BA is a potent HDAC inhibitor. To further demonstrate that 3-Cl BA acts as an HDAC inhibitor, we treated IECs with butyrate and 3-Cl BA (5 mM, 18 h) and assessed histone 3 (H3) acetylation levels as shown by the representative western blots in Fig. 2*E* and quantified in Fig. 2*F*. Similar to butyrate, 3-Cl BA significantly increased histone acetylation levels by protein, further suggesting that 3-Cl BA acts as an HDACi in IECs. Taken together, these data suggest that 3-Cl BA exhibits selective functions of native butyrate, namely HDACi, but does not stabilize HIF and is not used as a metabolic substrate. Such findings reveal that butyrate mimetics, such as 3-Cl BA, can be directed toward specific biological responses.

### 3-Cl BA regulates proteins involved in barrier function

Furthermore, we sought to compare native butyrate to 3-Cl BA on molecular endpoints in IEC. The intestinal epithelial barrier is regulated by the expression and localization of tight junctions (TJ) composed of several transmembrane and cytosolic proteins, including occludin (OCLN), claudins (e.g. CLDN2, CLDN4), zonula occludens, and cingulin (CGN).^40^ Based on the functional endpoints of promoting barrier formation and accelerated wound healing (Figure 1), we profiled the TJ regulation in several IEC lines. Interestingly, there was a remarkable transcriptional repression of *CLDN2* in Caco-2 cells exposed to butyrate and 3-Cl BA (both at 5 mM, 18 h) as well as the HDACi TSA (1 μM) as demonstrated by qPCR (Fig. 3*A*). CLDN2 is commonly referred as a “leaky claudin” that diminishes barrier integrity and a key component of a “leaky epithelia”.^41^ We extended these observations to assess if CLDN2 is repressed at a protein level. As shown in Figures 3*B* and 3*C*, there was a prominent loss of CLDN2 protein in Caco-2 cells exposed to butyrate or 3-Cl BA in PBS (5 mM, 24 h) as well as TSA (1 μM). A time course analysis revealed that butyrate and its mimicking compound 3-Cl BA decreased CLDN2 by as much as 80±3% after 48 h of exposure (Fig. 3*D* and *E*). These results were confirmed in a different IEC line (T84s), obtaining similar results in downregulation of *CLDN2* mRNA (same conditions as previously reported, Fig. 3*F*) and a significant suppression of protein (SI Appendix, Fig. 7*A-Bx*).

We extended these studies to profile other tight junction proteins in IEC. This analysis revealed a significant transcriptional upregulation of *CGN* in Caco-2 cells (SI Appendix, Fig. 7*C*), a protein localized to the cytoplasmic region of TJs and known to regulate CLDN2 expression.^42^ CLDN4 is a TJ-sealing claudin that correlates with tight epithelial tissues and competes with CLDN2 for residency within the TJ.^43^ We observed a significant upregulation of *CLDN4* exclusively in Caco-2 cells treated with 3-Cl BA (SI Appendix, Fig. 7*C*). Furthermore, synaptopodin (SYNPO) is localized to the actin cytoskeleton of intestinal epithelial TJ and is critical for barrier integrity and cell motility.^44^ We discovered that *SYNPO* is upregulated by both butyrate and the mimicking compound in Caco-2 cells (SI Appendix, Fig. 7*C*). In T84s treated with 3-Cl BA, we observed upregulation of *OCLN* (Fig. 3*F*), a TJ involved in barrier formation and that is markedly decreased in intestinal permeability disorders.^45^ Lastly, we transfected an OCLN-luciferase reporter construct in HeLa cells, treated the cells with butyrate or 3-Cl BA in PBS (5 mM, 24 h), and discovered a ∼6-7 fold increase in OCLN-luciferase reporter activity compared to vehicle, with no significant difference between butyrate and 3-Cl BA (Fig. 3*G*). Overall, these results indicated that both butyrate and its structurally-related mimic 3-Cl BA elicit a strong TJ phenotype that supports barrier integrity. In this regard, no differences were noted between native butyrate and 3-Cl BA.

### 3-Cl BA but not butyrate is protective in a mouse model of colitis

We extended our analysis of butyrate and 3-Cl BA to an in vivo model of colitis. To do this, we examined the impact of orally delivered butyrate and 3-Cl BA on colitis in wild-type (WT) C57BL/6 mice. Mice were treated with 2.5% dextran sulfate sodium (DSS), ±50 mM butyrate, or ±50 mM 3-Cl BA (both as sodium salts, pH 6-7) in drinking water. After 5 days of treatment, DSS was removed to allow mice to recover while continuing to consume H_2_O, butyrate, or 3-Cl BA solutions for an additional 2 days. As shown in Figure 4*A*, disease activity index (DAI) observations encompassing body weight, stool consistency, and bleeding show that all groups exhibited progressive disease while on DSS. Upon removing DSS, mice drinking 3-Cl BA recovered significantly faster than butyrate and water alone. Furthermore, mice exposed to 3-Cl BA significantly recovered initial weight after DSS removal compared to mice drinking butyrate or H_2_O (Fig. 4*B*). Upon sacrifice, 3-Cl BA treated mice exhibited a longer colon compared to the other groups (Fig. 4*C*). As shown in Figure 4*D*, intestinal barrier integrity was assessed at the end of the study via 4 kDa FITC-dextran oral gavages, showing that 3-Cl BA greatly contributed to promote a stronger intestinal barrier (29.0±8.3 μg/mL) compared to DSS only (143.4±43.2 μg/mL). We then assessed CLDN2 protein levels in distal colon. Consistent with our observations in vitro, CLDN2 protein was repressed both by butyrate and 3-Cl BA as demonstrated by the representative western blot (Fig. 4*E*) and relative CLDN2 densitometry (Fig. 4*F*). Lastly, we examined histologic features of disease in these cohorts. Strikingly, mice drinking 3-Cl BA exhibited far less tissue damage when compared to those exposed to butyrate or vehicle. Histology (Fig. 4*G-H*) revealed a significant protection afforded by 3-Cl BA compared to native butyrate. This analysis showed substantial loss of epithelial architecture, crypt damage, and inflammatory infiltrate both in butyrate/DSS and DSS-only treated mice. By contrast, histologic features of the 3-Cl BA/DSS-treated cohort more closely resembled healthy controls. These in vivo findings distinguish native butyrate and 3-Cl BA at the level of inflammatory protection.

**Figure 4.**
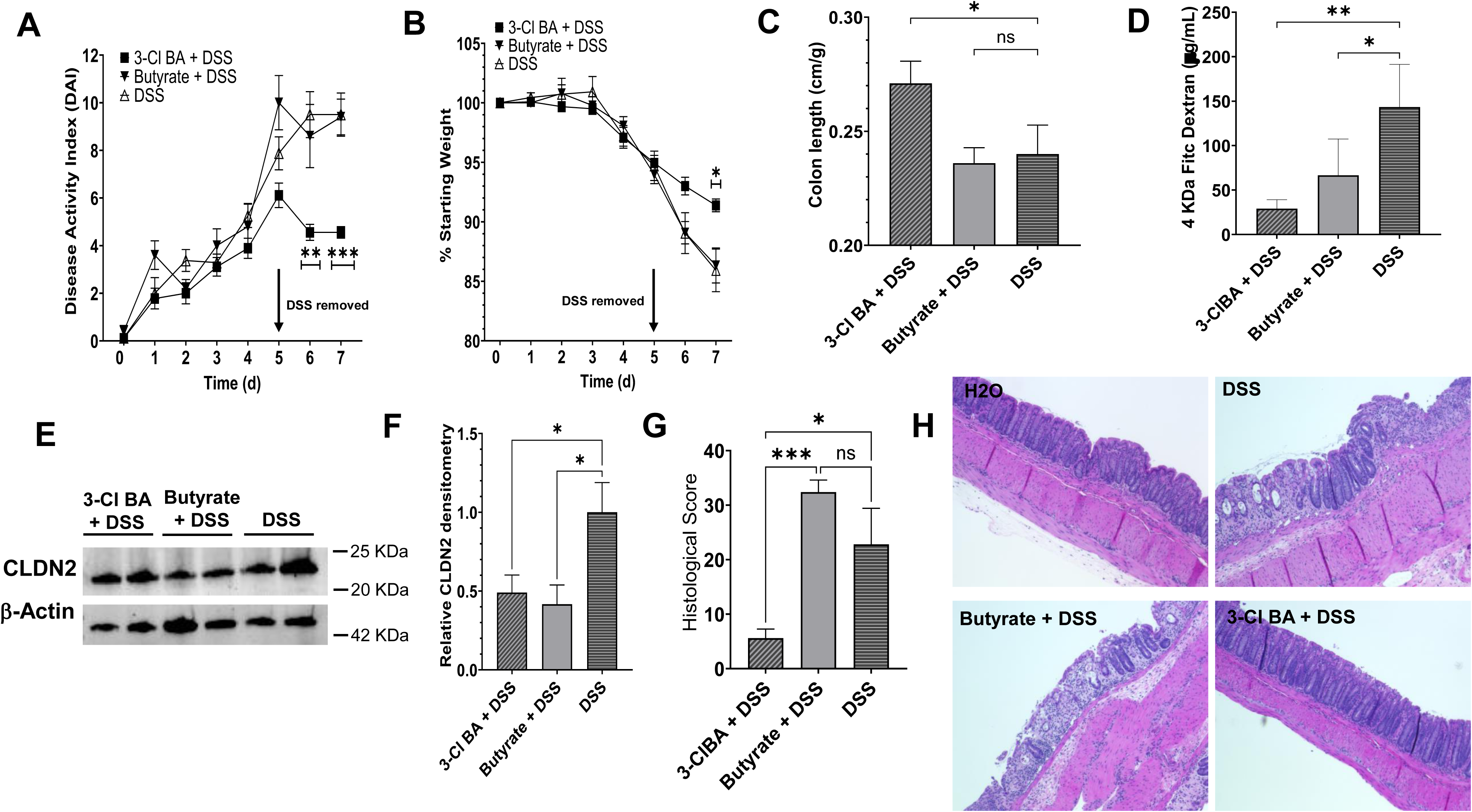
3-Cl BA but *not* butyrate is protective in a mouse model of colitis. C57BL/6 WT mice were exposed to 2.5% DSS +/-butyrate (50 mM) or 3-Cl BA (50 mM) (sodium salts) in their drinking water for 5 days. After day 5, DSS was removed but mice continued to drink butyrate or 3-Cl BA for two more days. (A) disease activity index score combining weight loss, stool consistency and bleeding and (B) %weight over time. (*n = 5-9*; error bars: SEM, \**p* < 0.05, \*\**p* < 0.01, \*\*\**p* < 0.001 by Two-way ANOVA with the Geisser-Greenhouse correction, Fisher’s multiple comparison). (C) Colon length to body mass ratios. (*n = 5-9*; error bars: SEM, \**p* < 0.05 by One-way ANOVA, Fisher’s multiple comparison). (D) Serum 4-KDa Fitc-dextran from mice subjected to DSS +/-butyrate or 3-Cl BA (*n = 4-5*, error bars: SEM, \**p* < 0.05, \*\**p* < 0.01 by One-way ANOVA, Fisher’s multiple comparison). (E) CLDN2 protein levels in distal colon tissue from mice exposed to different treatments and (F) quantification of CLDN2 protein using actin-normalized densitometry (*n = 4-5*, error bars: SEM, \**p* < 0.05 by One-way ANOVA, Fisher’s multiple comparison). (G) Histological scoring of colon tissue from mice exposed to DSS +/-butyrate or 3-Cl BA, maximum histological score of 40 (*n = 5*, error bars: SEM, \**p* < 0.05, \*\*\**p* < 0.001 by One-way ANOVA, Fisher’s multiple comparison). (H) Representative histology images showing healthy colon tissue (top left), the extent of colon tissue damage in mice subjected to DSS only (top right), DSS + butyrate (bottom left), and 3-Cl BA + DSS (bottom right).

### Comparison of butyrate and 3-Cl BA in mouse colonoids

To understand the dichotomy between butyrate and 3 Cl-BA in IEC cell lines and in vivo colitis model, we turned to colonoids derived from WT C57BL/6 mice. Stem-like mouse colonoids were plated on semipermeable membranes to allow for analysis of barrier formation. After 24 h, colonoids were allowed to differentiate in maturation media containing butyrate, 3-Cl BA (both at 1 mM), a bona-fide HIF stabilizer (PHD inhibitor IOX4, used at 10 μM) or vehicle (1% H_2_O). TEERs values were recorded every 24 h. As shown in Fig. 5*A*, this analysis revealed a striking difference in barrier formation between colonoids and transformed cells lines (T84 and Caco-2, see Fig. 1 and Fig. S1). Indeed, 3-Cl BA significantly enhanced barrier formation over vehicle while monolayers exposed to butyrate showed minimal barrier formation.

**Figure 5.**
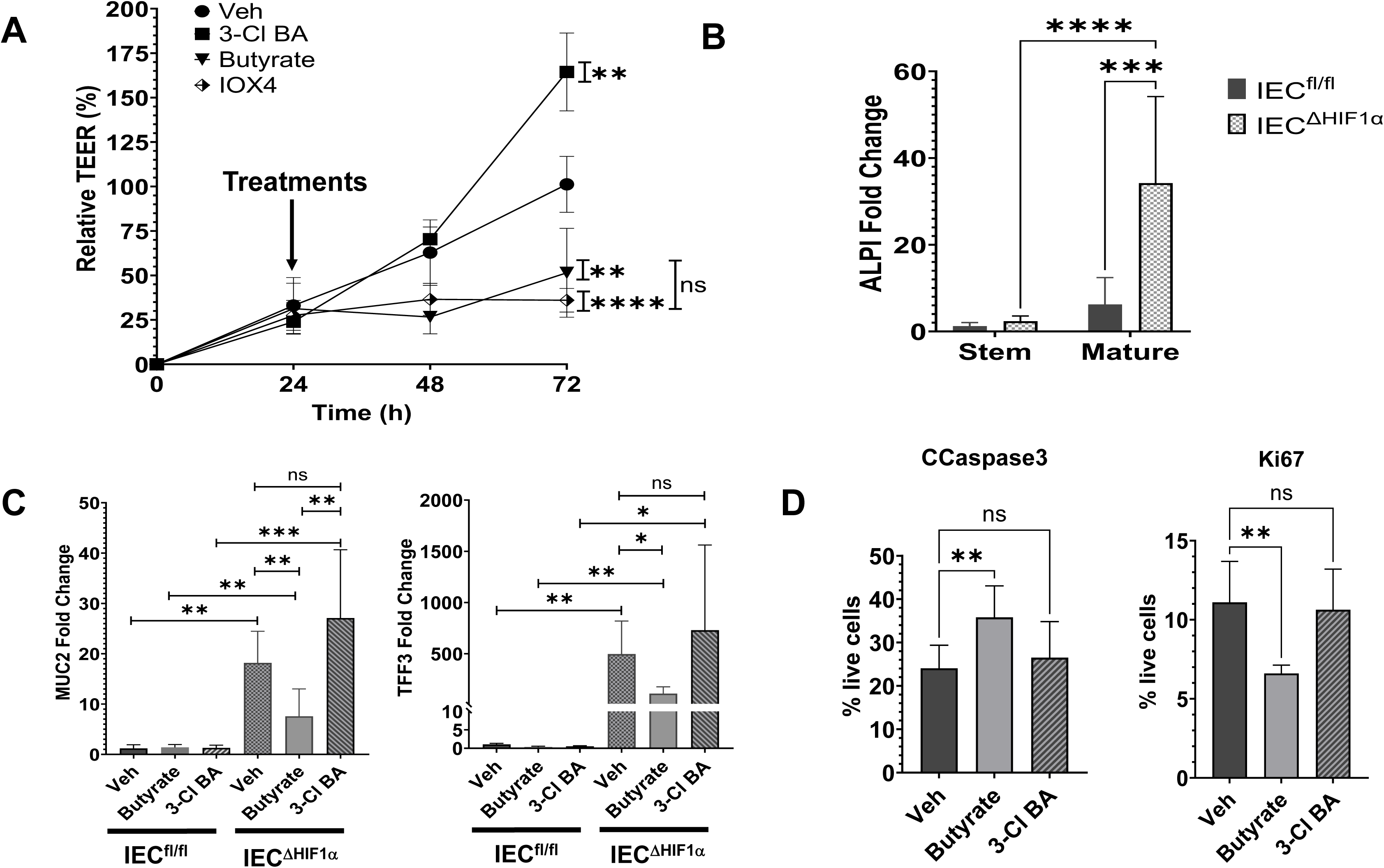
Influence of 3-Cl BA and butyrate in mouse-derived colonoids. (A) Stem colonoids (approximately 30,000 cells/well) were plated in semipermeable membranes and allowed to incubate for 24 h. Cells were then exposed to butyrate (1 mM), 3-Cl BA (1 mM), IOX4 (10 μM) or 1% H_2_O in maturation media. Barrier analysis by TEERs over time was conducted as previously described. Data is presented as relative TEER % normalized to vehicle treated cells at 72 h (100% TEERs) (*n = 8 inserts from 3 independent experiments*; error bars: SD, \*\**p* < 0.01, \*\*\*\**p* < 0.000 by Two-way ANOVA with the Geisser-Greenhouse correction. (B) *ALPI* mRNA expression in stem and mature IEC^fl/fl^ and IEC^ΔHIF-1α^ to evaluate absorptive intestinal epithelial cell differentiation. (*n = 6 wells from 3* independent experiments; error bars: SD, \*\*\*\**p* < 0.0001 by Two-way ANOVA, Tukey’s multiple comparison). (C) *MUC2* and *TFF3* mRNA changes in IEC^fl/fl^ and IEC^ΔHIF-1α^ exposed to maturation media with or without treatments for 48h to study differences in intestinal secretory cell differentiation. (*n = 6-9 wells from 3 independent experiments*; error bars: SD, \**p* < 0.05, \*\**p*< 0.01, \*\*\**p* < 0.001 by One-way ANOVA, Fisher’s multiple comparison). (D) Stem colonoids exposed to butyrate (1 mM), 3-Cl BA (1 mM) or 1% H_2_O were evaluated by flow cytometry for apoptosis and proliferation using CCaspase3 and Ki67 staining respectively. Frequency of Ki67 and CCaspase3-positive live cells is plotted. Data analyzed using FlowJo software (*n = 6-7 wells from 2 independent experiments*; error bars: SD, \*\**p* < 0.01, One-way ANOVA, Fisher’s multiple comparison.

One of the prominent differences discovered between butyrate and 3-Cl BA was the stabilization of HIF (see Fig. 2A). Interestingly, like butyrate, the potent HIF-stabilizing compound IOX4 significantly inhibited barrier formation in mouse colonoids (Fig. 5*A*), suggesting that HIF stabilization in stem cells disrupts barrier formation and likely explains our observed differences between stem cells and differentiated transformed cell lines. To further investigate the role that HIF plays in IEC differentiation, we isolated colonoids from mice with intestinal epithelial specific deletion of HIF-1α (IEC^ΔHIF-1α^) and Villin Cre negative as a control (IEC^fl/fl^). We studied differences in absorptive intestinal epithelial cell differentiation by comparing IECs kept in stem media (stem) to IECs incubated in maturation media (mature) for 48 h. As observed in Fig. 5B, using ALPI as a differentiation marker, we observed that the deletion of HIF-1α significantly accelerates maturation. Furthermore, we exposed stem-like IEC^fl/fl^ and IEC^ΔHIF-1α^ to butyrate or 3-Cl BA (both at 1 mM) in maturation media over 48 h. Fig. 5C demonstrates significant differences in secretory cell differentiation when HIF-1α is lacking as observed by a potent upregulation of *MUC2* and *TFF3*. Furthermore, we discovered that butyrate, but not 3-Cl BA, significantly reduced the induction of these differentiation markers in our IEC^ΔHIF-1α^ cell line. This could be attributed to the fact that butyrate stabilizes HIF-2α and may play a role in repressing differentiation. These results support the important role that HIF plays in IEC stem cells suggesting that it inhibits or reduces differentiation.

Lastly, WT colonoids were exposed to butyrate or 3-Cl BA (1 mM) in stem media over 24 h to elucidate their effects on cell death, proliferation and growth in stem cells. Flow cytometry studies revealed that only butyrate significantly promotes IEC apoptosis as observed by increased cleaved caspase3 levels (CCaspase3) and inhibits proliferation as indicated by reduced Ki67 (Fig. 5*D*). Thus, the dichotomy between our observations in transformed cell lines and in vivo are likely explained by differential responses of IEC stem cells to butyrate and 3-Cl BA where HIF stabilization likely plays a major role.

## Discussion

Microbially-derived metabolite mimicry is a relatively novel and underexplored concept that could accelerate the discovery of new drugs from their naturally-derived moieties. SCFAs, for example, exhibit multiple functions in a location-dependent manner. While it is well-accepted that butyrate functions as a preferred energy source for colonocytes, it also orchestrates transcriptional control of a multitude of genes as an HDAC inhibitor and through the stabilization of HIF in the mucosa^46^. Given the diverse functions of this relatively simple metabolite, we aimed to determine the existence of butyrate-mimicking compounds that might regulate specific butyrate biological pathways as a potential rescue mechanism in disease-associated dysbiosis. Here we report the identification of a novel butyrate-mimicking compound, namely 3-Cl BA, that regulates intestinal barrier function and mucosal wound healing selectively through HDACi control of genes involved in TJ formation.

Butyrate is known to exhibit an overall positive influence on mucosal homeostasis^37^. As the primary energy source for IECs, butyrate contributes to maintaining a hypoxic microenviroment by promoting mitochondrial oxidative phosphorylation and increasing oxygen consumption.^23,24^ In health, this hypoxic environment, termed “physiologic hypoxia”, is paramount for epithelial integrity and homeostasis. By contrast, dysbiosis of butyrate-producing microbes strongly correlates with the severity of IBD and other mucosal inflammatory diseases^5^. We assessed the impact of butyrate-mimicking compounds and elected to use epithelial barrier formation as a model biological function. Initially, we screened a small library of butyrate mimetics for their influence on epithelial barrier integrity using immortalized cancer cell lines *in vitro*.^47–49^. We discovered that, similar to native butyrate, 3-Cl BA selectively increased barrier formation in IECs. Interestingly, other butyrate-mimicking compounds (select examples described here) had no barrier-enhancing activity, including the closely related structural isomer 4-chlorobutyrate (4-Cl BA). Furthermore, a possible hydrolysis byproduct from 3-Cl BA, 3-hydroxybutyrate, did not exhibit barrier-promoting activity *in vitro*. In parallel, wound healing assays revealed that 3-Cl BA accelerated wound healing compared to other butyrate-mimicking compounds, including 4-Cl BA. Further studies revealed that both butyrate and 3-Cl BA promoted a strong tight junction profile *in vitro* by potently downregulating a leaky claudin and upregulating other key components in the TJs.

These promising results led us to investigate the underlying mechanism for this 3-Cl BA selectivity. For these purposes, we profiled three of the major functions of butyrate in the colon, including as a fuel for oxidative phosphorylation, as an HDACi and as a HIF stabilizing agent^46^. This analysis revealed that 3-Cl BA is not readily metabolized by oxidative phosphorylation and results in no appreciable HIF stabilization. HDACi assays, however, revealed that similar to butyrate, 3-Cl BA is a potent HDACi. This observation is mostly consistent with studies by Verma, et. al who screened thirty-six SCFA-like compounds (3-Cl BA was not included in the study) to define the most potent HDACi activity^50^. These studies revealed that native butyrate showed the most HDACi activity followed closely by molecules modified at the carbon 3 position. Likewise, they demonstrated that butyrate inhibited HDACs by binding to the Zn^2+^ in the catalytic site of HDACs. In this regard, the installation of a halogen functional group in the third position, such as in 3-Cl BA, could facilitate its position in the binding site resulting in strong HDACi capabilities. Taken together, this evidence suggests that 3-Cl BA is a strong candidate as a selective HDACi in colonocytes.

Our *in vivo* results comparing administration of native butyrate and the 3-Cl BA mimetic during DSS colitis were surprising. Although all cohorts of mice became ill within 5 days of DSS exposure regardless of treatment, mice exposed to 3-Cl BA recovered rapidly, with only minimal histologic evidence of colitis by day 7. In contrast, animals administered butyrate showed strong clinical signs of colitis with histologic evidence of profound inflammation. While surprising, these results are not completely unexpected. The use of native butyrate for therapeutic purposes is, at best, controversial. Studies in both murine models and in human colitis have revealed that administration of butyrate can be beneficial, neutral and even detrimental^51^. It is thought that several individual factors may contribute to these outcomes, including dosage, uptake and metabolism of butyrate during active inflammation. An additional consideration is the impact of native butyrate on ISCs. Since stem cell proliferation is essential for effective wound healing, it is notable that butyrate has been reported to inhibit ISC proliferation^30^. Based on this evidence, we extended our *in vivo* experiments to compare butyrate or 3-Cl BA *ex vivo* on stem-like mouse colonoids. These studies confirmed previous reports that butyrate inhibits ISC proliferation while the mimetic 3-Cl BA showed a more favorable response in enhancing the formation of differentiated epithelial monolayers.

A plausible explanation for the observed differences between butyrate and 3-Cl BA on colonic stem cell proliferation may lie in the architecture of the intestinal crypt. It has been demonstrated, for instance, that a natural metabolite gradient stemming from the top of the villus to the bottom of the crypt results in the rapid consumption of butyrate as fuel by differentiated colonocytes, thereby minimizing exposure of the crypt to butyrate.^30^ In crypt-less organisms (e.g. zebra fish) or intestinal injury (e.g. DSS colitis), this gradient is disrupted by loss of villus architecture and ISCs are unnaturally exposed to metabolites such as butyrate. Along these same lines, the intestinal crypt also exhibits a prominent oxygen gradient (physiologic hypoxia). Oxygen concentrations drop precipitously along the crypt-villus axis from the well-oxygenated crypt floor with constant blood flow (pO_2_ ∼85) to the top of the villus where the microbiota and metabolites reside in a hypoxic environment (pO_2_ <10).^52^ Recently, it was demonstrated that the constitutive expression of specifically HIF-1α impairs ISC activity. The authors demonstrated that a proliferative phenotype is significantly amplified in mice with specific deletion of HIF-1α. Furthermore, organoid formation was more effective in HIF-1α deficient ISC’s compared to controls.^53^ These results suggest that HIF spatially influences the intestinal epithelial dynamics and that ISC’s at the bottom of the crypt are maladapted to hypoxic conditions and the regulatory influences of HIF. Based on the observation that butyrate, but not 3-Cl BA, stabilizes HIF, we assessed the influence of HIF on stem-like colonoid barrier formation. Using a pharmacological HIF stabilizing drug IOX4 under normoxic conditions, it was determined that HIF stabilization phenocopied butyrate by inhibiting stem cell proliferation and blocking barrier formation of colonoids. Furthermore, we also confirmed that the specific deletion of HIF-1α in mouse colonoids potently accelerates differentiation of ISCs to absorptive or secretory IECs. We discovered that butyrate significantly inhibits ISC differentiation in IEC^ΔHIF-1α^ possibly through HIF-2α stabilization. Furthermore, we observed that butyrate inhibits proliferation of WT ISCs and promotes cell death. Butyrate inhibition of stem cell proliferation has also been attributed to the transcription factor FOXO3^30^. It is notable that HIF-1α can positively and negatively regulate FOXO3^54^. Thus, we propose that stem cell HIF stabilization by butyrate, but not 3-Cl BA, could explain the differences in ISC proliferation observed in our studies.

Taken together, we demonstrate that it is possible to identify microbial-derived metabolite-mimicking compounds with selective functions. Furthermore, we highlight the importance of a dual metabolic/oxygen gradient within the intestinal crypt and the differences in spatial effects of potential therapeutics. Using this rationale, we identified 3-Cl BA, a molecule with selective HDACi properties that lacks other biological properties of native butyrate. While we do not know the exact details of the protection afforded by 3-Cl BA in active inflammation, we infer that by avoiding the detrimental impact that native butyrate plays on stem-cell proliferation and differentiation, 3-Cl BA enhances wound healing responses *in vivo*. This work provides a foundation for the development of metabolite-mimicking therapeutics as an approach to discover bioactive small molecules for the treatment of diseases where dysbiosis predominates, such as IBD.

## Materials and Methods

### Cell culture, treatments, and chemical reagents

T84 cells (ATCC #CCL-248) were cultured in DMEM/F-12 1:1 (Thermo Fisher Scientific) containing Pen/Strep, GlutaMAX, and 10% (v/v) heat-inactivated bovine calf serum (BCS, Hyclone). Clones of Caco-2 cells (Caco-2, ATCC #CRL2102)^55^ were grown in IMDM (Corning) containing Pen/Strep and Glutamax + 10% BCS. All the cell lines were maintained at 37°C and 5% CO2 in a humidified incubator. Cells were passaged approximately every 7 days with trypsin (Thermo Fisher Scientific). Cells were plated on 6-well or 24-well plates with or without semipermeable inserts, as required. Sodium butyrate, 3-chloro butyric acid, 4-chloro butyric acid, gamma-amino butyric acid, crotonic acid, 3-hydroxy butyric acid, and 4-KDa Fitc-dextran were acquired from Sigma Aldrich, Trichostatin A (Selleck Chemicals), and PBS (Fischer Scientific). Unless otherwise noted, cells were exposed to a final concentration of 5 mM BA or BA derivatives, previously neutralized with NaOH (sodium salts), by adding appropriate volumes of 50 mM solutions prepared in PBS directly in the media and PBS was used as vehicle treatment control.

### Protein analysis and immunoblotting

For western blotting, cells were immediately placed on ice after treatments and cell lysates were prepared by scraping in a freshly prepared cocktail of ice-cold 1X Laemmli buffer (Bio-Rad) containing 1X HALT protease inhibitor (Thermo Scientific), 5 mM EDTA, and 100 mM DTT (Sigma Aldrich) in H_2_O, (450 μL per well in a 6 well-plate). The obtained lysate solutions were sonicated until their viscosity were like that of water. Mouse tissue samples were lysed using radio-immunoprecipitation (500 μl, RIPA) lysis buffer containing 1X HALT protease inhibitor and 5 mM EDTA, sonicated and centrifuged. The protein concentration was determined by BCA protein assay (Thermo Fisher Scientific). Iced-cold tissue lysates (63 μl) were mixed with the previously described Laemmli cocktail (37 μl) and equal concentration proteins were resolved. SDS-PAGE samples were run on Mini-PROTEAN TGX precast gels 10% (Bio-Rad) and transferred to 0.2 μm PVDF membranes using a Bio-Rad Transblot Turbo system. Blots were blocked for 1 h at room temperature using blocking buffer 5% milk (Bio-Rad) in Tris-buffered saline (TBST) (25 mM Tris-HCl, 150 mM NaCl, 0.1% Tween-20) and then incubated overnight in primary antibody diluted in 5% milk in TBST. The primary antibodies used were anti-β-actin (Abcam #ab8227, 1:5000), anti-HIF-1α (BD Biosciences #610959, 1:1000), anti-HIF-2α (Novus Bio #NB100-122, 1:1000), anti-acetyl histone 3 (Active Motif #39139, 1:1000), anti-histone 3 (Millipore Sigma #06-942, 1:1000), and anti-CLDN2 (Fischer Scientific #51-6100, 1:1000). Blots were then washed with TBST (3X, 10 min) and incubated for 1 hour with their corresponding HRP-conjugated secondary antibody diluted in 5% milk (1/10000). Blots were washed thoroughly with TBST, developed using Clarity MAX ECL reagent (Bio-Rad), and imaged using a Bio-Rad ChemiDoc MP imager. Densitometry was measured using ImageJ, comparing the bands of interest to the Actin bands.

### RNA isolation, cDNA synthesis and qPCR

RNA was isolated from cells using TRIzol reagent (Thermo Scientific) according to the manufacturer’s instructions. Isolated RNA was then reverse transcribed to cDNA using the iScript Supermix reagent (Bio-Rad), and analyzed using Power SYBR Green Master Mix reagent (Thermo Scientific) in a Quant Studio 5 Applied Biosystems Real-Time PCR (Thermo Fisher Scientific). Transcript quantities were calculated using an on-plate standard curve and were normalized to that of β-actin. The following primer sequences were used for real-time PCR analysis: *hactb*, forward – 5’ – CATGTACGTTGCTATCCAGGC - 3’, reverse, 5’ - CTCCTTAATGTCACGCACGAT – 3’; *hcldn2*, forward- 5’ – GCCTCTGGATGGAATGTGCC - 3’ reverse, 5’ – GCTACCGCCACTCTGTCTTTG – 3’; *hocln*, Forward – 5’ - GTCATCCACGAGGCGAAGTTAAT- 3’, reverse, 5’ – ACAAGCGGTTTTATCCAGAGTC – 3’; *hcldn4*, forward – 5 ‘- GGGGCAAGTGTACCAACTG -3’, reverse, 5’ – GACACCGGCACTATCACCA – 3’; h*cgn*, forward - 5’ -TGGAAAGCTACTCCGTTCCCA -3’, reverse, 5’ – AGCAGTGTCAATGGTGCTACC – 3’; *hsynpo*, forward - 5’ - ATGGAGGGGTACTCAGAGGAG - 3’, reverse, 5’ – CTCTCGGTTTTGGGACAGGTG – 3’; *mactb*, forward - 5’ – AACCCTAAGGCCAACCGTGAA – 3’, reverse, 5 – TCACGCACGATTTCCCTCTCA – 3’; *mlgr5*, forward – 5’ – CCTACTCGAAGACTTACCCAGT – 3’, reverse, 5’ – GCATTGGGGTGAATGATAGCA – 3’; *malpi*, forward – 5’ – ATGATGCCAACCGAAACCCC – 3’, reverse, 5 – GCGTGTCCTTCTCATTGGTAA – 3’; *mtff3*, forward – 5 – TAATGCTGTTGGTGGTCCTG – 3’, reverse – 5 – CAGCCACGGTTGTTACACTG – 3’; *mmuc2*, forward – 5’ – TGCCCACACACTTTGGAGAG – 3’, reverse – 5’ – CCTCACATGTGGTCTGGTTG – 3’.

### Assessment of epithelial barrier function

Epithelial barrier was measured by transepithelial electrical resistance (TEER) and FITC dextran flux assay. TEER was measured by growing IECs on permeable membranes, 0.4 μm pore, 24 well plate using an epithelial voltmeter (model EVOM2, World Precision Instruments). All treatments (5 mM unless otherwise noted) were applied 24 h after plating the cells and TEER was monitored every 24 h. Raw measurements were converted to Ohm*cm^2^. 4 KDa FITC-dextran flux was measured when cells were at maximal barrier as measured by TEER. Media was replaced with 1 mL of Hanks+ buffer in the bottom well and the top well was filled with 80 μl of 1.25 μg/μl of FITC-dextran 4KDa in Hanks+ Buffer. Samples were obtained from the lower chamber every 30 minutes for 2 hours and measured relative to a standard curve. Flux was measured as μg/min/cm^2^. C57BL/6 colonoid TEER experiments were conducted by dissociating colonoids in Trypsin-EDTA (Fisher Scientific) by vigorous pipetting (colonoid development described below). Cells were strained through a 70 μm filter, counted and then plated in equal number per condition (approximately 30,000/well) on collagen coating solution (Sigma-Aldrich, 125–50) pretreated transwells. To collagen coat transwells, 0.4 μm pore, 0.33cm^2^ transwells were incubated with 100 μl of collagen coating solution for 12–24 h prior to removal of excess by suction. Cells were grown in L-WRN cell conditioned enteroid media for 24 h, followed by maturation media (L-WRN media diluted 1:5 with T84 media) containing the corresponding treatments: BA and 3-Cl BA (1 mM), IOX4 (10 μM), vehicle = H_2_O. TEERs were obtained every 24 h. Fitc-dextran assay and flux calculation were done as described above without modifications.

### Wound healing assay

T84 cells were plated at 35,000 cells/well on a 96-well ImageLock plate (Essen Bioscience Inc.) and incubated until a confluent cell monolayer formed (∼48 h). Precise and reproducible wounds were made in all wells with a WoundMaker (Essen Bioscience Inc.). After wounding, media was aspirated from each well, and each well was gently washed with PBS before 100 μL of control media or media containing treatments (BA or BA derivatives, all at 5 mM) were added. Initial images were taken immediately after wounding at 10X using the IncuCyte live-cell imaging (Essen Bioscience Inc.), and then every 2 h over the course of 36 h. Relative wound closure% was quantified for every image using the relative wound density metric, a measure of cell density in the wound area relative to the cell density outside of the wound area.

### Real-time oxygen consumption

A SensorDish Reader from Applikon Biotechnology and oxodish plates (Precision Sensing) were used as previously described.^56^ Briefly, Caco-2 monolayers were grown to confluency on 0.33 cm^2^ transwell inserts with 0.4 μm pore size (Corning). Once fully confluent as observed by maximum barrier formation, the medium was aspirated, and inserts were transferred to a 24-well oxodish plate with an oxygen sensor at the bottom center of each well. One milliliter of medium was added basolaterally and 250 μl to the apical side and allowed to equilibrate in an incubator for 1 h at 37° C. Corresponding treatments were added directly to the media, and the oxodish was immediately placed on the sensordish reader on a rotating platform at 37° C. The plate was not sealed to allow for reoxygenation of the media. The oxygen percentage in the media was continuously measured at 1-min intervals. O_2_ consumption rates were calculated by linear regression of individual plots.

### HDAC inhibition assay

HDAC activity/inhibition were measured using Epigentek kits (P-4034-96) following their protocol without modifications. Briefly, Caco-2 cells were grown to confluency in 6-well plates. Then, nuclear extracts were obtained using a NE-PER extraction kit (Thermo Fisher Scientific) following their protocol without modifications. The obtained protein was quantified using BCA protein assays as described by regular protocols. Nuclear extracts (∼4 ug per well) were added to the HDAC assay plate and incubated with treatments: BA and 3-Cl BA (5 mM), TSA (1 μM), vehicle = PBS and the provided assay buffer. Wells were then washed with provided washing buffer, followed by the addition of capture antibody. Wells were washed again, and detection antibody was added. Lastly, a color developing solution was added and the absorbance was measured. HDAC activity calculation was performed as instructed by the company.

### Transfection

an occludin-luciferase reporter plasmid and empty vector control (Switch-gear Genomics) were transfected into HeLa cells using Lipofectamine 3000 transfection reagent (Invitrogen) following the manufacturer’s recommended protocol and as previously reported.^57^ The day following transfection, cells were treated with 5 mM BA or 3-Cl BA in PBS (24 h). Cells were lysed the day after treatment and luciferase was measured using a dual-luciferase reporter assay (Promega #E1960) and the results were normalized to protein by BCA.

### WT and HIF-1α KO colonoid development and treatments

WT colonoids were developed using colon samples from C57BL/6 wildtype mice. Intestinal epithelial specific deletion of HIF-1α in mice was generated by crossing HIF-1α floxed mice^58^ (Jax stock #007561) with Villin Cre positive mice^59^ (JAX stock #004586). Colonic tissue derived organoids were isolated from WT mice or mice with an intestinal epithelial specific deletion of HIF-1α or Villin Cre negative (C57BL/6) control, as previously described^38,60,61^. To summarize, approximately 2-3 cm of proximal colonic tissue was harvested from the appropriate mouse, washed, and opened lengthwise. The colonic tissue was chemically digested using the Lamina Propria Dissociation Kit (Miltenyi Biotech, 130-097-410) and subsequently mechanically digested on the intestine setting using a GentleMACs Dissociator and GentleMACs C tubes (Miltenyi Biotech, 103-293-237). Digested samples were incubated at 37°C until epithelial crypt dissociation, subsequently strained through a 70 μM strainer, washed, and plated in Matrigel (Corning, 354234). The colonoids were passaged 2-3 times weekly and split at a ratio of 1:1-1:3. Colonoids were grown in stem media consisting of L-WRN conditioned media, supplemented with EGF 50 ng/mL (R&D, 2028), nicotinamide 10 mM (R&D, 4106), A83-01 500 nM (R&D 2939), SB202190 10 μM (R&D 1264), CHIR99021 3 μM (R&D, 4423), thiazovivin 4 μM (R&D, 3845), SB431542 4 μM (R&D, 1614), gastrin I 10 nM (R&D, 3006), and Y-27632 10 μM (Tocris, 1254), supplied by the University of Colorado Organoid and Tissue Modeling shared resource, and 50% Intesticult Organoid Grown Media when necessary (Mouse) (StemCell Technologies, 06005). For maturation experiments, IEC^fl/fl^ or IEC^ΔHIF-1α^ were plated in Matrigel as spots on tissue culture plates and incubated for 24 h in stem media. Colonoids were then switched to maturation media prepared by a 1:5 dilution of stem media:T84 cultured media and incubated for 48 h with or without treatments (1 mM butyrate or 3-Cl BA, Veh = H_2_O). Treatments were applied at the same time as maturation media. Colonoids were collected in Trizol for qPCR analysis and processed the same as other cell lines (T84s, etc) for downstream analysis.

### Flow cytometry

C57BL/6 WT colonoids were cultured as previously described in Matrigel domes in 6-well plates for 48 h using stem media. After this time, they were treated with 1 mM BA or 3-Cl BA in H_2_O for 18 h. Isolation of mouse colonoids and analysis were performed as previously described.^62^ Briefly, to generate single-cell suspensions, cultures were incubated with ice-cold PBS to loosen the Matrigel, followed by mechanical dissociation through repeated pipetting. Dissociated colonoids were collected in ice-cold PBS and centrifuged at 150 × g for 10 minutes at 4°C. After washing to remove residual Matrigel, colonoids were incubated with pre-warmed TrypLE Express for 3 minutes at 37°C, then further dissociated by gentle pipetting. TrypLE was neutralized with MACS buffer supplemented with 5% FBS, and cells were centrifuged at 400 × g for 5 minutes at 4°C. The final pellet was resuspended in MACS buffer + 5% FBS at an approximate concentration of 10 million cells/mL.

For flow cytometry, cells were first stained with Live/Dead Blue viability dye (1:1000 dilution) for 20 minutes on ice in the dark. Samples were washed with MACS buffer + 5% FBS and centrifuged at 400 × g for 5 minutes at 4°C. For intracellular staining, cells were fixed and permeabilized using the fixation/permeabilization solution from BD Biosciences, following the manufacturer’s instructions. After permeabilization for 1 hour at 4°C in the dark, cells were stained with anti-Ki-67 (PE, 1:100) and anti-Cleaved Caspase-3 (Alexa Fluor 647, 1:50) in permeabilization buffer. Following staining, cells were washed and resuspended in IC Fixation Buffer (BD Biosciences) according to the manufacturer’s protocol. Samples were acquired on a Cytek Aurora 5-laser spectral flow cytometer. Data were analyzed using FlowJo software, version 10.10.0.

### Mouse studies

C57BL/6 mice were purchased from Jackson Laboratories. Animals were maintained and bred in standard housing conditions under 24 h/day, 7 days/week veterinary care at the University of Colorado Anschutz Medical Campus (AMC) animal facility. Animal procedures and care were reviewed and approved by the Institutional Animal Care and Use Committee of AMC.

### DSS-induced colitis

8- to 13-week-old mice of similar weight and matched by gender were used in DSS studies. Mice were switched to water bottles in their cages to allow them to become familiarized with this drinking system 24 h before the experiments began. On day 0, the DSS group received drinking water with 2.5% DSS (MW ∼40,000; Chem Impex), the BA + DSS group received 2.5% DSS + 50 mM BA (sodium salt) in water, and the 3-Cl BA + DSS group received 2.5% DSS + 50 mM 3-Cl BA (sodium salt) in water. Treatments were given for 5 days, and fresh solutions were prepared and replaced every other day. After this time, mice were allowed to recover for 2 days by removing DSS but still treated with either 50 mM BA or 3-Cl BA in their drinking water. A disease activity index (DAI) score was assessed daily to evaluate the development of colitis based on the parameters of weight loss compared to initial weight, stool consistency, and rectal bleeding. Scores were defined as weight loss: 0 (0%), 1 (1-5%), 2 (5-10%), 3 (11-20%), and 4 (>20%); stool consistency: 0 (well-formed pellets), 2 (pasty, semi-formed pellets), and 4 (liquid stools); and rectal bleeding: 0 (no blood), 2 (hemoccult positive), and 4 (gross bleeding). Maximum DAI possible was 12. Colon lengths were measured at time of sacrifice, and distal colon tissue collected for histology and protein analyses.

### Colon Permeability

Mice were administered 100 μL of 100 mg/mL 4-kDa FITC-dextran (Sigma-Aldrich) by oral gavage, 2 h later blood was collected by heart puncture immediately after sacrifice.

### Histological Scoring

Distal colon samples were fixed in methacarn (methanol:chloroform:acetic acid, 60:30:10) before preparation for histological analysis and staining with hematoxylin and eosin. Colon tissue from a healthy mouse without any treatments was used as reference and shown in the H_2_O histology image. All histological quantitation was performed blinded and scored by a pathologist. Each sample was assessed for three independent parameters including severity of inflammation (0-3: none, slight, moderate, severe), depth of injury (0-3: none, mucosal, mucosal and submucosal, transmural), and amount of crypt damage (0-4: none, basal 1/3 damaged, basal 2/3 damaged, only surface epithelium intact, entire crypt and epithelium lost). Each independent parameter score was then multiplied by a factor reflecting the percentage of tissue involved (x1: 0-25%, x2: 26-50%, x3: 51-75%, x4: 76- 100%) and then totaled. Maximum histological score possible was 40.^63^

### Nuclear magnetic resonance spectroscopy

^1^H and ^13^C NMR were recorded on a Bruker Avance III HD (400 MHz) NMR instrument. Residual solvent signal was used as internal standard.

### Statistical and graphical presentation of data

Statistical analysis and figure generation were performed using GraphPad Prism 10. Statistical analyses were performed using either One-way or Two-way ANOVA with post-hoc corrections, as indicated in the figures. A *p* value of less than 0.05 was considered significant.

## Abbreviations

HIFs: hypoxia-inducible-factors
HDAC: histone deacetylase
HDACi: histone deacetylase inhibitor
ISCs: intestinal stem cells
3-Cl BA: 3-chlorobutyrate
DSS: dextran sulfate sodium
SCFAs: short-chain fatty acids
IBD: inflammatory bowel disease
IECs: intestinal epithelial cells
TEERs: transepithelial electrical resistance
4-Cl BA: 4-chlorobutyrate
3-OH: 3-hydroxybutyrate
GABA: γ-aminobutyrate
FITC-dextran: fluorescein isothiocyanate-dextran
NMR: nuclear magnetic resonance
PBS: phosphate-buffered saline
TJ: tight junctions
H3: histone 3
OCLN: occludin
CLDN2: claudin-2
CLDN4: claudin-4
CGN: cingulin
SYNPO: synaptopodin
DAI: disease activity index
WT: wild-type
MUC2: mucin-2
TFF3: trefoil factor-3
BCS: bovine calf serum
CCaspase3: cleaved caspase3

## Author contributions

A.O. and S.P.C. conceptualized the idea and designed the project; A.O., J.A.C., J.Y.K, R.H.C, B.D.G, R.L.R, F.M., K.Y.V.S., L.Z. performed research; J.L.M.D, P.R., I.C., C.H.T.H, G.B., J.C.O., A.S.D. were instrumental in providing analytical tools, reagents, interpretation of results, and contributing relevant scientific expertise to the project; A.O. and S.P.C. analyzed data and wrote the manuscript.

## Data availability statement

All data generated in this study are included in this article and the supplementary information or are available from the corresponding author upon reasonable request.

---

**Figure S1.**
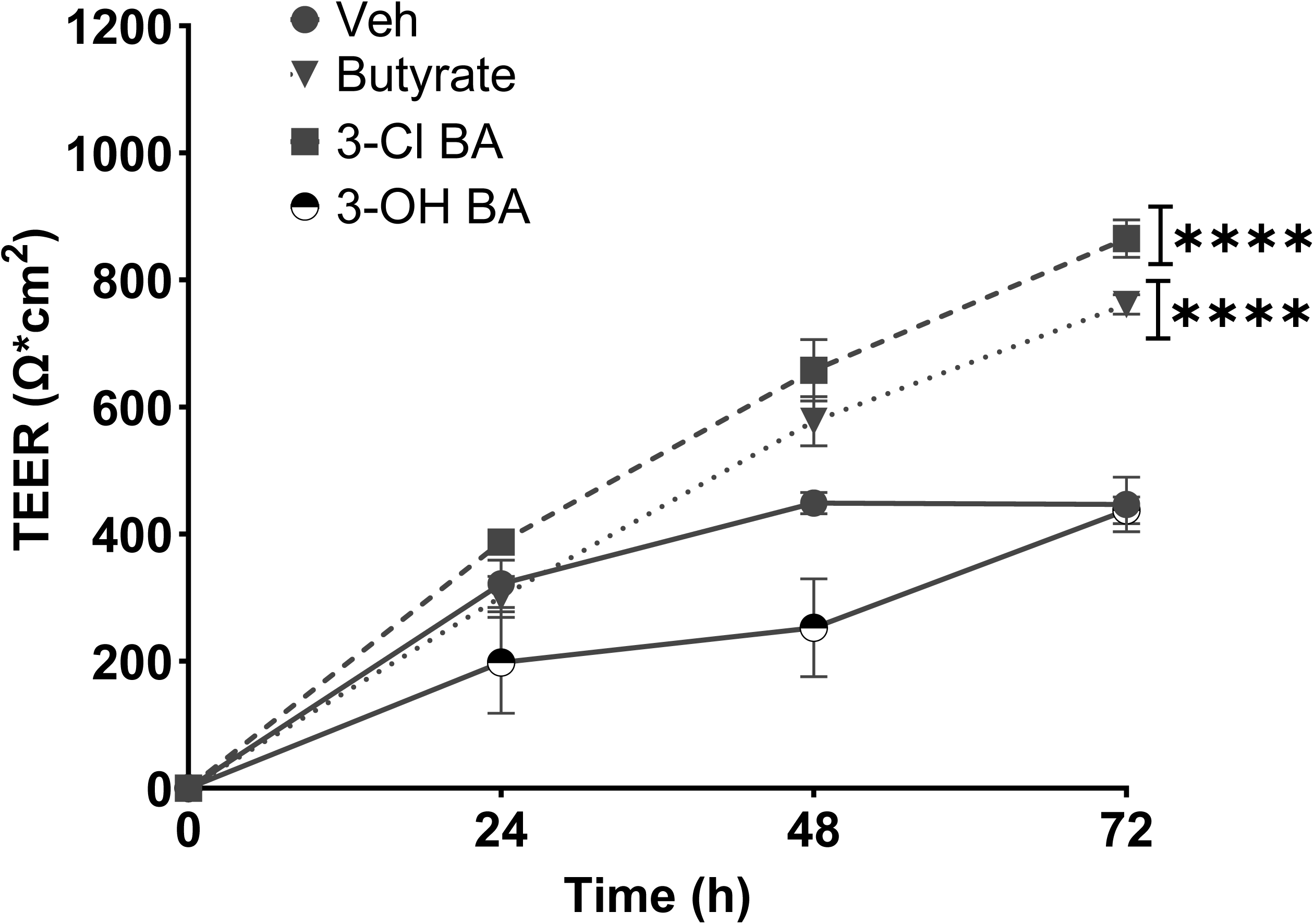
Influence of butyrate and butyrate-mimicking compounds in Caco-2 IEC barrier formation. Epithelial barrier formation over time in monolayers of Caco-2 cells exposed to butyrate, 3-Cl BA, and 3-OH BA (all at 5 mM, Veh = PBS). (*n = 3;* error bars: SEM, \*\*\*\**p* < 0.0001 by Two-way ANOVA with the Geisser-Greenhouse correction, Dunnett’s multiple comparisons test).

**Figure S2.**
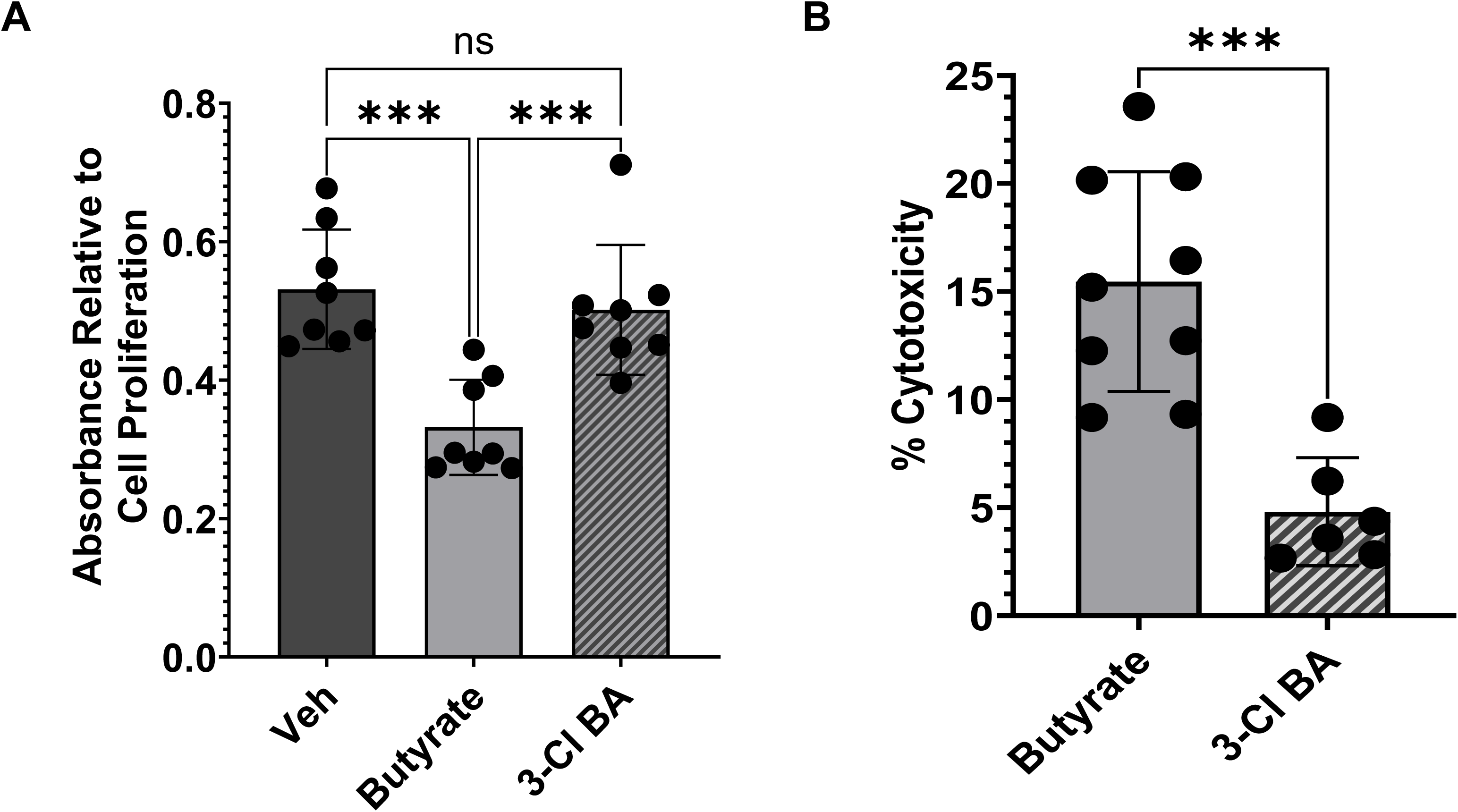
Proliferation and cytotoxicity studies in T84s exposed to butyrate or 3-Cl BA. (A) Cell proliferation in T84 cells exposed to butyrate or 3-Cl BA over 24 h. Data presented as absorbance relative to cell proliferation (*n = 8* wells from 3 independent experiments; error bars: SD, \*\*\**p* < 0.001 by One-way ANOVA, Fisher’s multiple comparison). (B) Percent cytotoxicity observed after exposing T84 cells to 5 mM butyrate or 3-Cl BA (24 h). (*n = 6-9* wells from 3 independent experiments; error bars: SD, \*\*\**p* < 0.001 by One-way ANOVA, Fisher’s multiple comparison).

**Figure S3.**
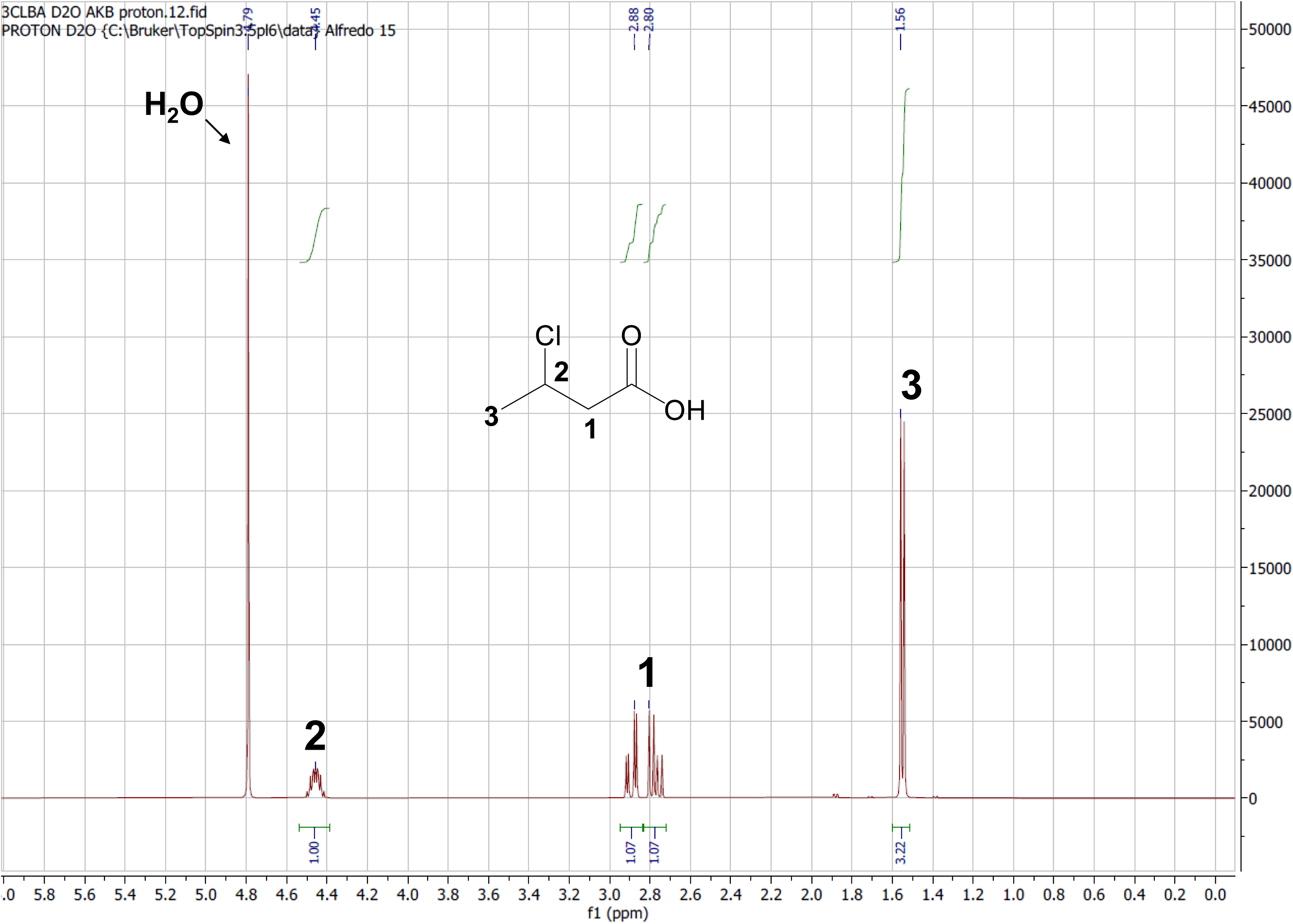
^1^H NMR spectrum of 3-Cl BA in deuterated water (D_2_O). ^1^H NMR (Bruker NMR 400 MHz, 295 K, D_2_O) δ 4.79 (H_2_O residual peak), 4.45 (m, 1 H, **2**), 2.88-2.80 (dd, 2 H, **1**), 1.56 (d, 3 H, **3**). The proton in the COOH group is not visible due to a rapid exchange with deuterium from D_2_O causing the signal to be undetectable.

**Figure S4.**
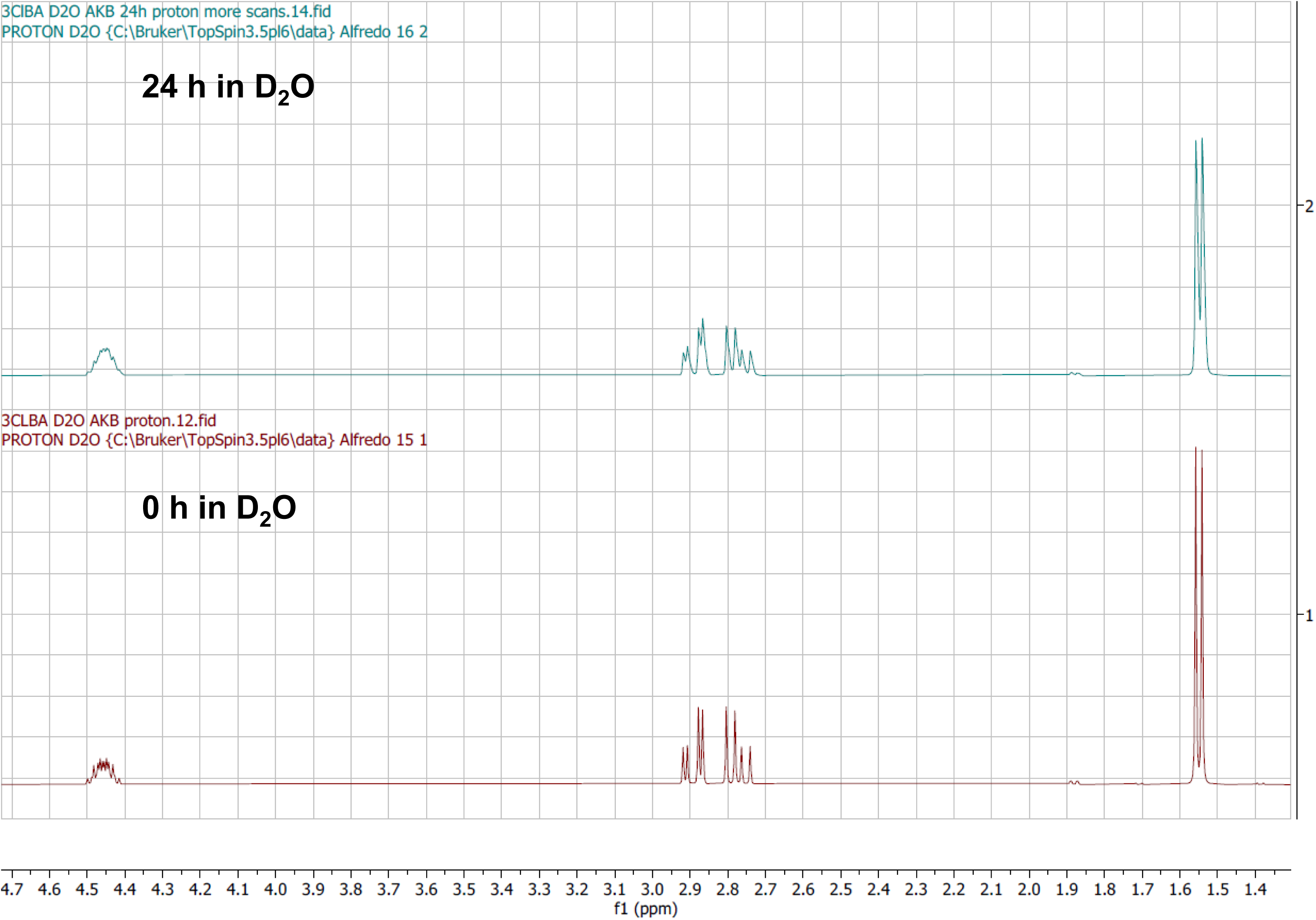
Comparison of 3-Cl BA ^1^H NMR spectra over 24 h to establish stability of compound in water. 3-Cl BA (25 mgs) was dissolved in ∼0.4 mL of D_2_O and a ^1^H NMR was immediately taken. The compound was allowed to incubate in deuterated water at 37 °C for 24 h followed by another ^1^H NMR experiment. As seen here, there is no change in the integrity of the compound over this time.

**Figure S5.**
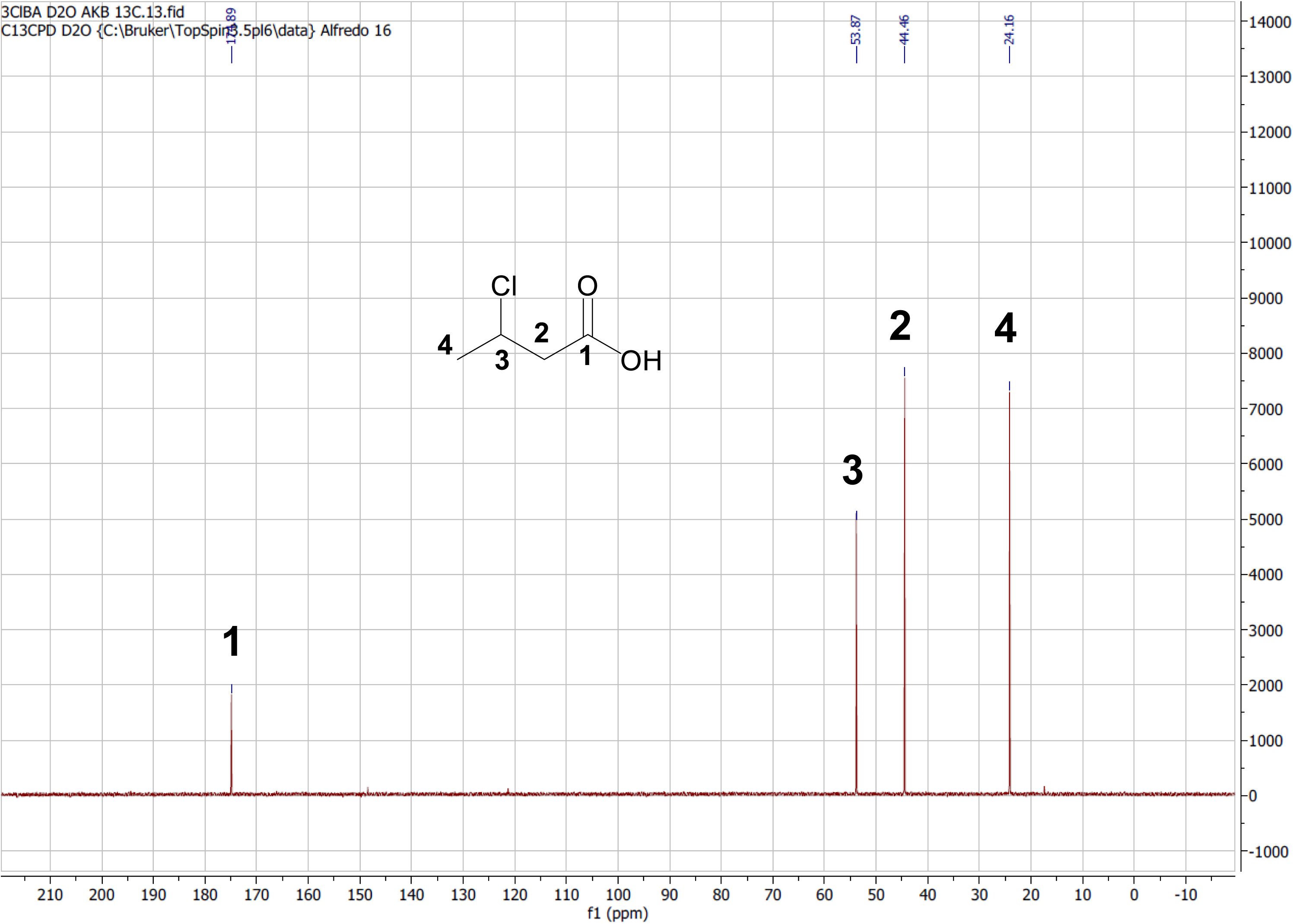
^13^C NMR spectrum of 3-Cl BA in D_2_O. ^13^C NMR (Bruker NMR 100 MHz, 295 K, D_2_O) δ 174.9 (**1**), 53.9 (**3**), 44.5 (**2**), 24.2 (**4**).

**Figure S6.**
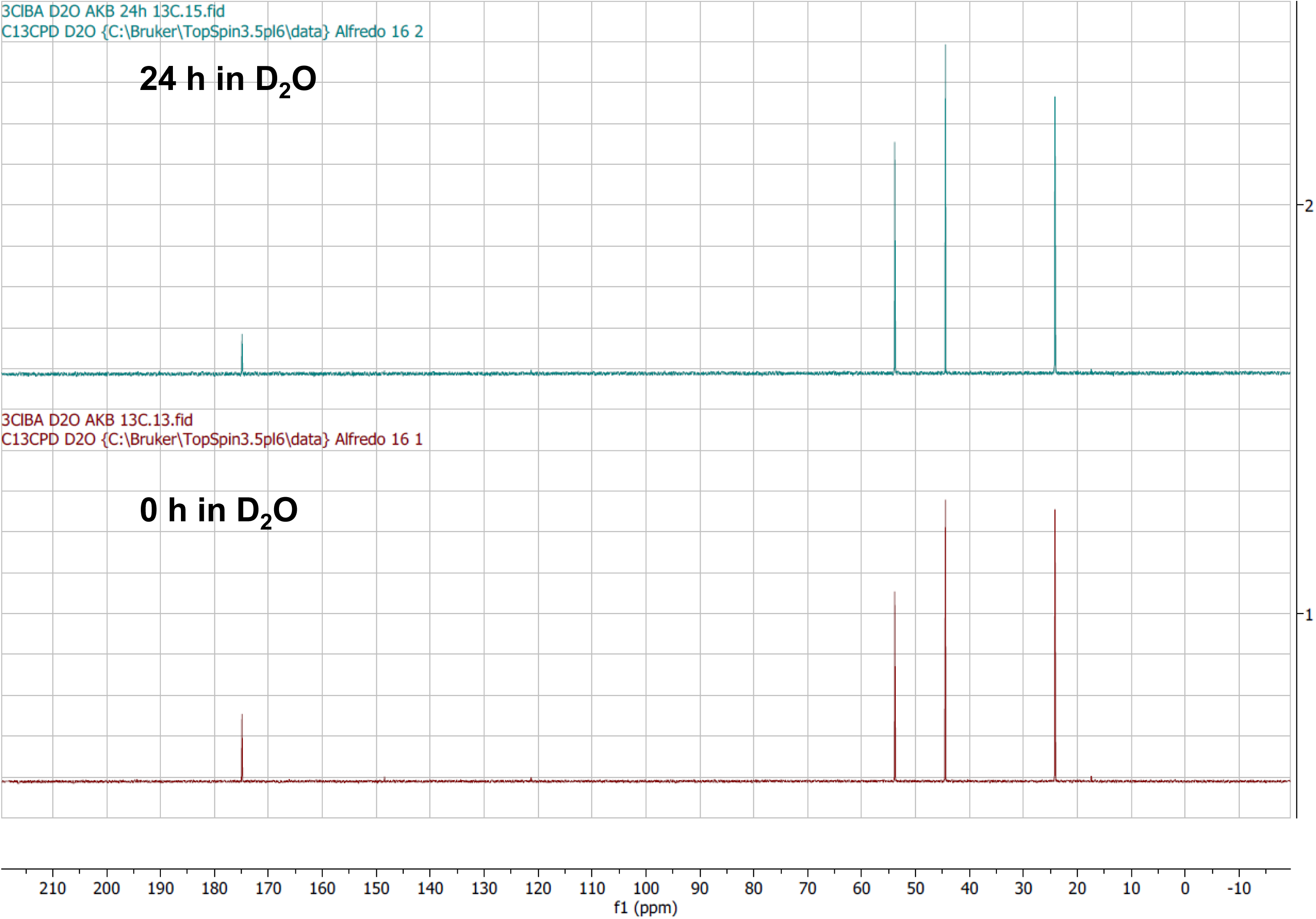
Comparison of 3-Cl BA ^13^C NMR spectra over 24. **h.** Following the same conditions described before, ^13^C NMR spectra were compared demonstrating no change in the integrity of the compound when dissolved in water over 24 h.

**Figure S7.**
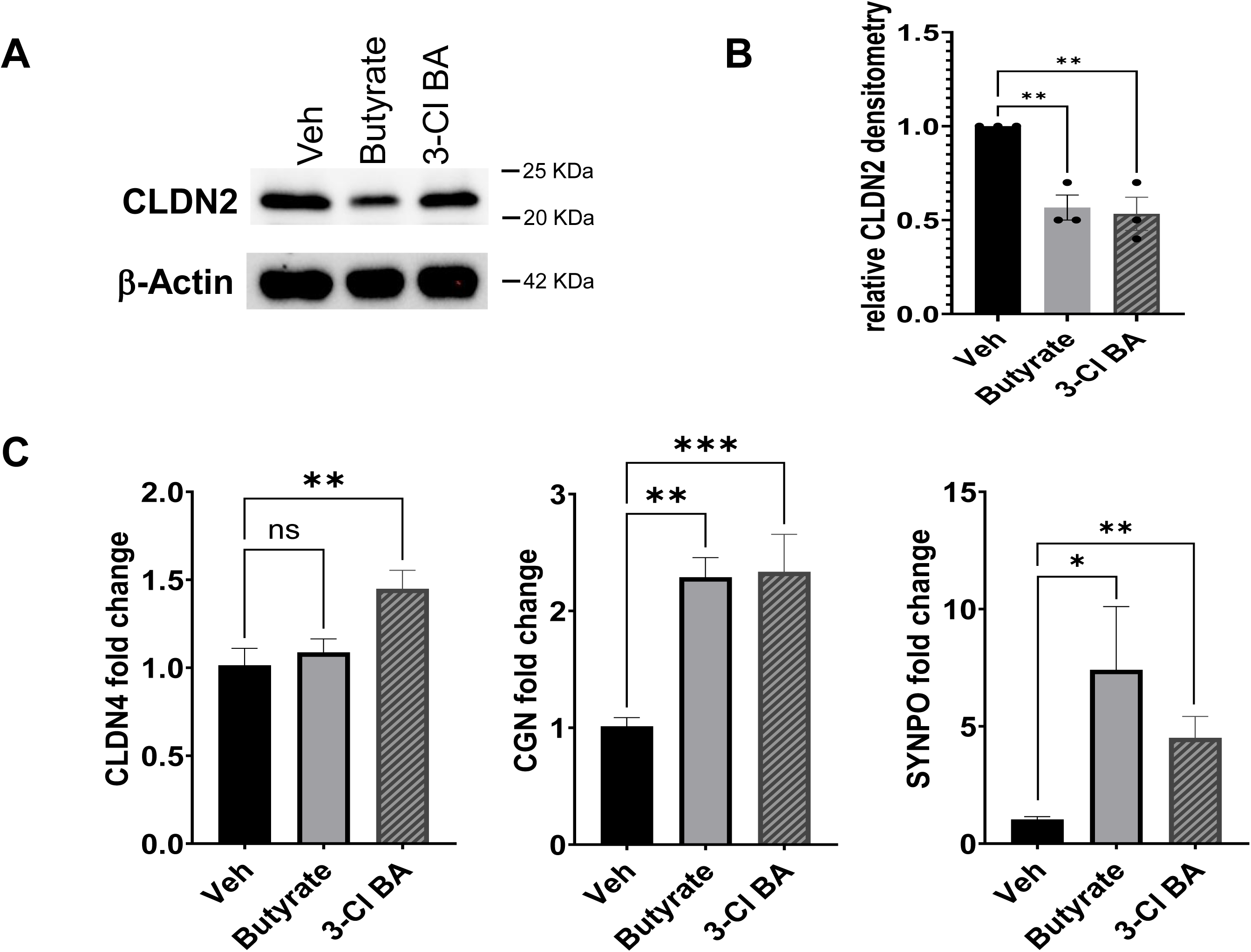
Tight junction profiles in IECs exposed to butyrate or 3-Cl BA. (A) CLDN2 protein levels in T84 cells treated with 5 mM butyrate or 3-Cl BA, Veh = PBS. Cells were exposed to treatments for 24 h. (B) Quantification of CLDN2 protein was performed using actin-normalized densitometry. (*n = 3*; error bars: SEM, \*\**p* < 0.01 by One-way ANOVA, Fisher’s multiple comparison). (C) *CLDN4,* CGN, and SYNPO mRNA expression in Caco-2 cells exposed to 5 mM butyrate, 3-Cl BA or Veh = PBS for 18 h. (*n = 3*; error bars: SEM, \**p* < 0.05, \*\**p* < 0.01, \*\*\**p* < 0.001 by One-way ANOVA, Fisher’s multiple comparison).

